# Genotypic and phenotypic evidence indicates the introduction of two distinct forms of a non-native species (*Gekko gecko*) to Florida, USA

**DOI:** 10.1101/2023.02.07.527561

**Authors:** Thomas W. Fieldsend, Herbert Rösler, Kenneth L. Krysko, Madison E.A. Harman, Stephen Mahony, Timothy M. Collins

## Abstract

The red-spotted tokay gecko *Gekko gecko* (Linnaeus, 1758) is a widely distributed Asian gecko that has established outside of its native range in Florida, USA. This study used genotypic and phenotypic data to determine whether multiple, distinct forms of red-spotted tokay gecko are present in Florida, specifically the morphologically distinct “*G. g. gecko*” and “Form B”. Two discrete mitochondrial clades (ND2) were identified in Florida tokay geckos, the native range distributions of which were found to correspond closely with the native ranges of *G. g. gecko* (the Malay Archipelago) and Form B (mainland Asia), respectively. Furthermore, each mitochondrial clade was non-randomly associated with a separate nuclear (RAG-1) clade, supporting the existence of two discrete genotypic lineages, representative of *G. g. gecko* and Form B. Both lineages were detected in Florida, and population-level morphological structure was non-randomly associated with genotype, confirming a genotype-phenotype link. Multiple lines of evidence thus indicate the introduction of both forms of tokay gecko to Florida, where hybridisation likely also occurs. The link between morphological Form B and a unique genotypic lineage also indicates the existence of a tokay gecko taxon distinct from the recognised subspecies *G. g. azhari* Mertens, 1955 and *G. g. gecko* (Linnaeus, 1758).

## Introduction

The tokay gecko *sensu lato* is a large (up to 181 mm snout-vent length; Rösler 2005), colourful gecko with a vast native range spanning from Nepal in the northwest to the Aru Islands (eastern Indonesia) in the southeast (Rösler et al. 2011). Once common, the tokay gecko has declined in many parts of its native range (Caillabet 2013, Weterings et al. 2019). Millions of tokay geckos are harvested annually, both for the Asian traditional medicine trade—where they are used in the treatment of ailments including AIDS and asthma (Bauer 2009, Nijman et al. 2012a, Nijman et al. 2012b, Caillabet 2013, Nijman and Shepherd 2015)—and for the international pet trade (Nijman et al. 2012b, Casey et al. 2015). Despite these native-range declines, non-native tokay gecko populations are now established in Curaçao (Behm et al. 2019), Florida, USA (King and Krakauer 1966, Krysko et al. 2019), Martinique (Henderson et al. 1993, Breuil 2009), Mexico (Villalobos-Juárez et al. 2022), Puerto Rico (Romero Cordero 2021), and likely Belize (J Meerman, pers. comm.) and Hong Kong (Dufour et al. 2022), due both to deliberate introductions (e.g., King and Krakauer 1966, Breuil 2009) and accidental escapes (e.g., Behm et al. 2019). As a result of multiple intentional releases in Florida from the 1960s onwards (King and Krakauer 1966, Krysko et al. 2019), tokay geckos have established throughout much of the state (Krysko et al. 2019), and have been reported from as far south as Key West (Meshaka et al. 2004) and as far north as Leon County (Means 1996), which borders Georgia.

*Gekko gecko* (including *Gekko verticillatus* Laurenti, 1768 as its junior synonym) is the type species of the genus *Gekko* (Wood Jr et al. 2020). Despite being heavily studied (Reese et al. 2004, Rösler et al. 2011), the taxonomy of the tokay gecko *sensu lato* is unsettled (Kongbuntad et al. 2016, Saijuntha et al. 2019). Rösler (2005) recognised four morphologically distinct forms of tokay gecko: *G. g. azhari* Mertens, 1955, distributed in Bangladesh and possibly adjacent parts of India; *G. g. gecko* (Linnaeus, 1758), from Philippines, Brunei, Indonesia, the Malaysian islands of Sarawak and Sabah, and possibly parts of mainland southeastern Asia (e.g., peninsular Malaysia); “Form A” from northern Vietnam, southern China, and Taiwan; and “Form B”, with a widespread distribution including Nepal, Laos, Cambodia, and Thailand, and possibly also northern India, southern Vietnam, the Andaman Islands, and peninsular Malaysia. Although Rösler (2001) noted that *G. g. azhari* shares some meristic and morphometric features with the *G. smithii* complex (recently reviewed in Grismer et al. 2022), subsequent examination of *Gekko* populations located close to the *G. g. azhari* type locality (Chittagong Hills Tract, Bangladesh) leaves little doubt as to their closer morphological affinity to the *G. gecko* complex (pers. obs.; Mahony and Reza 2008, Mahony et al. 2009). Form A was elevated to species status by Rösler et al. (2011) on morphological grounds, under the revalidated name *G. reevesii* (Gray, 1831). The authors restricted the native range of *G. reevesii* to China and northern Vietnam, noting that the tokay geckos present in Taiwan were probably introduced from mainland China. They also noted that the name *G. reevesii* corresponds to the “black” tokay—distinguished by its black coloured dorsal spots (e.g., Figure 10 in Rösler et al. 2011)—as distinct from the “red” tokay, which possesses red or orange dorsal spots (Das 2002). Notwithstanding the wide variation in colouration exhibited by tokay geckos, these two primary colour morphs have long been recognised (Rösler et al. 2011), and are generally referred to as the “black-spotted” and “red-spotted” tokay, respectively (e.g., Peng et al. 2010, Wang et al. 2012). The tokay gecko *sensu lato* is thus currently polytypic, comprising the black-spotted tokay gecko *G. reevesii*, and the red-spotted tokay gecko *G. gecko*, to which belong the subspecies *G. g. gecko* and *G. g. azhari*, as well as the morphologically distinct Form B, which is presently of unassigned taxonomic rank.

Saijuntha et al. (2019) performed a phylogenetic analysis of 166 native-range tokay geckos using the NADH dehydrogenase 2 (ND2) mitochondrial (mtDNA) gene, and reported the existence of five distinct mtDNA clades (i.e., Clades A–E in Figure 4 of Saijuntha et al. 2019). mtDNA Clade A was the only clade recovered from specimens from the islands of southeastern Asia, and mtDNA Clade A haplotypes were also detected on mainland southeastern Asia in Cambodia, peninsular Malaysia, northeastern Thailand, and northern Vietnam. mtDNA Clade B was found to be widely distributed across mainland Asia, including parts of Myanmar and Laos and much of Thailand. mtDNA Clade C was represented by a single specimen from Magway, which is located in Myanmar’s Central Dry Zone (Herridge et al. 2019). mtDNA Clade D was found to occur sympatrically with mtDNA Clade B in parts of northeastern Thailand and central Laos, while mtDNA Clade E was only detected in northern Thailand. No *G. g. azhari* sequences were analysed, while purported *G. reevesii* sequences were polyphyletic and present in both mtDNA Clade A and mtDNA Clade B.

Fieldsend et al. (2021b) sequenced 751 bp of the ND2 gene of 34 tokay geckos collected from introduced populations in Florida, and detected haplotypes belonging to both mtDNA Clade A and mtDNA Clade B. On the basis of the large pairwise genetic distances (up to 11.77%) observed between mtDNA Clade A and mtDNA Clade B haplotypes in Florida, the authors speculated that these mtDNA clades might represent distinct forms of red-spotted tokay gecko, and thus the introduction of multiple tokay gecko forms to Florida. Furthermore, the authors noted a close phylogeographical correspondence between these two mtDNA clades and two of the morphologically distinct tokay gecko forms described in Rösler (2005). mtDNA Clade A is ubiquitous on the islands of southeastern Asia, and is present in relatively restricted areas on the mainland, including peninsular Malaysia; its distribution thus closely matches that of *G. g. gecko sensu* Rösler (2005). Moreover, the distribution of mtDNA Clade B accords well with the hypothesised distribution of Form B *sensu* Rösler (2005) (see above). Rösler (2005) also noted that a contact zone between *G. g. gecko* and Form B might exist on the Malay Peninsula, and the presence of both mtDNA Clade A (e.g., Kedah, Malaysia, GenBank accession number JN019051; Rösler et al. 2011) and mtDNA Clade B (e.g., Takua Pa, Thailand, GenBank accession number MK117103; Saijuntha et al. 2019) haplotypes on the Malay Peninsula (see Fig. 4 of Saijuntha et al. 2019) provides further evidence that these two morphologically distinct tokay gecko forms may also be genetically distinct.

One aim of this study was to establish whether multiple distinct forms of red-spotted tokay gecko—specifically, *G. g. gecko* and Form B—have indeed been introduced to Florida, as hypothesised by Fieldsend et al. (2021b). This is important partly because hybridisation or admixture between distinct forms could lead to elevated population-level genetic diversity (e.g., Kolbe et al. 2004, Kolbe et al. 2007) or ‘hybrid vigour’ (Facon et al. 2005), and thus increase the invasive potential (Facon et al. 2005, Lavergne and Molofsky 2007, Crawford and Whitney 2010, Wagner et al. 2017, Smyser et al. 2020) of what may already be an invasive taxon (Fieldsend et al. 2021a, Fieldsend et al. 2021b). Moreover, distinct forms of tokay gecko could differ markedly in life history traits relevant to establishment success in Florida (e.g., bioclimatic niche; Zhang et al. 2014), and thus pose differing degrees of threat to Florida’s native biodiversity.

A second aim of this study was to determine whether Florida tokay gecko populations could be used to better clarify the taxonomy of the red-spotted tokay gecko. Kongbuntad et al. (2016) and Saijuntha et al. (2019) both postulate that the red-spotted tokay gecko may represent an unresolved species complex; if so, then the chance to examine multiple red-spotted tokay gecko forms in Florida would present an unusual opportunity to use an introduction event to study an unresolved species complex (e.g., Wegener et al. 2019).

## Methods

Forty-five tokay geckos were collected in southern Florida between 18 January 2019 and 08 February 2021 and were accessioned into the Florida Museum of Natural History (FLMNH), Gainesville, Florida, USA (see Appendix S1). Geckos were euthanised via intracoelomic injection of MS222 (tricaine methanesulfonate), transported on dry ice, and stored at -80 °C as per Fieldsend et al. (2021b), under Florida International University IACUC protocol # IACUC-17-019. Additionally, four tokay gecko specimens were borrowed from the Field Museum of Natural History (FMNH 180852 and FMNH 180849, Nakhon Ratchasima Province, Thailand; FMNH 266245, Zambales Province, Philippines; FMNH 236071, Romblon Province, Philippines), with tissue samples also received for the two Philippines specimens. Given their native-range provenance, these four tokay geckos were expected to represent *G. g. gecko* (Philippines) and Form B (Thailand) (Rösler 2005, Rösler et al. 2011, Saijuntha et al. 2019).

Genetic analysis of specimens broadly followed Fieldsend et al. (2021b). mtDNA sequences were generated for a part of the ND2 mitochondrial coding gene, along with a full tRNA^Trp^ sequence and part of tRNA^Ala^ (“ND2”). Sequence data were also generated for the recombination activating gene 1 (RAG-1) nuclear (nDNA) coding gene. Total DNA was extracted using cetyltrimethylammonium bromide (CTAB) (Saghai-Maroof et al. 1984), and double-stranded DNA was amplified via polymerase chain reaction (PCR) in 50 μl reactions as detailed in Fieldsend et al. (2021b). PCR conditions for double-stranded DNA amplification of ND2 were as per Fieldsend et al. (2021b). RAG-1 amplification was achieved with the primers from Wang et al. (2013) (F: CCAGAGGAAGTTCAGCAGTGTC; R: GCTTCCAACTCATCAGCTTGTC) using an initial hot-start step of 10 min at 57 °C, followed by 3 min at 95 °C, followed by 36 cycles of 35 s of 95 °C denaturation / 45 s of 58 °C annealing / 150 s of 72 °C extension, followed by 60 s at 49 °C, followed by a final 15 min extension period at 72 °C. ExoSap-IT (Thermo Fisher Scientific, Waltham, MA, USA) was used to clean double-stranded PCR products, which were then cycle-sequenced using using BigDye™ Terminator v3.1 Cycle Sequencing Kit (Thermo Fisher Scientific, Waltham, MA, USA) as per Fieldsend et al. (2021b). Finally, an Applied Biosystems 3130XL Genetic Analyzer was used to read sequences on both strands. Chromatograms were visually examined for heterozygous peaks. No heterozygous peaks were found in ND2 sequences. RAG-1 sequences with only one heterozygous peak (*n* = 7) were taken to represent alleles differing by one base pair, and were coded at this position with the appropriate IUPAC code. For reads with > 1 heterozygous peak, attempts were made to clone into plasmid vectors. However, in several instances sequencing of multiple Colony Forming Units resulted in ≥ 3 putative alleles being returned, potentially due to polyploidy (Evans et al. 2005), the presence of pseudogenes (Mighell et al. 2000), contamination, or cloning error. Consequently, sequences obtained via cloning were considered unreliable and were omitted from subsequent analyses. Sequences were successfully obtained for Florida tokay geckos for ND2 (*n* = 45) and RAG-1 (*n* = 26). All 26 RAG-1 and 11 ND2 sequences were wholly novel, while the remaining 34 ND2 sequences were extensions (1090 bp) of the 751 bp sequences originally published in Fieldsend et al. (2021b). Novel ND2 and RAG-1 sequences were also generated for the two Philippines specimens. Sequences generated as part of this study were deposited into GenBank (Sayers et al. 2019) under accession numbers OL631485-OL631559.

Sequence alignments (Appendix S2) were generated using MUSCLE (Edgar 2004) in MEGA X (Kumar et al. 2018), and comprised sequences both from this study (see above) and retrieved from GenBank. GenBank sequences were predominantly generated by Rösler et al. (2011), and were used because they represent georeferenced native-range specimens for which both ND2 and RAG-1 have been sequenced. Additional sequences from Macey et al. (1999), Townsend et al. (2004), Gamble et al. (2008), Jackman et al. (2008), and Wang et al. (2013) were also included in the alignments.

Three separate sequence alignments were created: 1) ND2 sequences only (1090 bp) (*n* = 54); 2) RAG-1 sequences only (553 bp) (*n* = 35); and 3), an additional, full data alignment created by concatenating 1090 bp of ND2 and 553 bp of RAG-1 (1643 bp) for all specimens (Florida and native-range) for which both sequences were available (*n* = 34) (Appendix S2). As it could not be determined whether the ND2 (GenBank accession number AF114249; Macey et al. 1999) and RAG-1 (GenBank accession number AY662625; Townsend et al. 2004) sequences from Phuket, Thailand correspond to the same specimen, they were not concatenated. All sequence alignments contained a *G. chinensis* sequence to serve as an outgroup, because it is the most-closely related species to *G. gecko* (Rösler et al. 2011, Wood Jr et al. 2020) for which suitable ND2 and RAG-1 sequence data were available that did not cluster within *G. gecko* for RAG-1 in preliminary analyses (e.g., *G. smithii*, unpublished data). The inclusion of *G. chinensis* sequences meant that the number of tokay gecko sequences in each alignment was *n*-1.

Bayesian phylogenetic analyses were performed in MrBayes (v.3.2.7; Ronquist et al. 2012) using a mixed-model approach, which results in the Markov chain sampling over the space of all possible reversible substitution models (Ronquist et al. 2012). In each case, Bayesian posterior probabilities (BPPs) were determined by running one million generations of Metropolis coupled Markov chain Monte Carlo [(MC)^3^], with two simultaneous runs of four chains, each starting with a random tree, with sampling occurring every 100 generations. Preliminary runs were conducted to ensure stationarity of the dataset, with the potential scale reduction factor approaching one for all parameters, and the standard deviation of split frequencies **≤** 0.01. One quarter of trees were discarded as burnin, with the remaining three-quarters used to construct the 50% majority-rule consensus tree and estimate BPPs. A partitioned analysis (Ronquist et al. 2012) was performed on the concatenated alignment, with the ND2 and RAG-1 sequence data treated as separate partitions, and parameters unlinked across the partitions.

Morphological analysis of tokay geckos was based predominantly on a subset of the morphological traits described in Rösler (2005) and Rösler et al. (2011), which have proven useful in differentiating between different forms of tokay gecko. Sex of Florida specimens was confirmed by necropsy, as the external morphological traits generally used for sexing tokay geckos (i.e., cloacal spurs, pre-cloacal pores, and post-cloacal sacs) can be ambiguous (pers. obs.). The native-range specimens were not sexed, as necropsies were not permitted by the loaning organisation. To control for any potential ontogenetic shifts in morphological traits, only specimens with a snout-vent length (SVL) ≥ 110 mm were analysed, as this is the approximate size at which tokay geckos of both sexes mature in Florida (Meshaka et al. 2004). Likewise, traits based on ratios (e.g., head length / head height; Rösler et al. 2011) were not utilised, as tokay geckos continue to grow beyond sexual maturity (Kurniati and Phadmacanty 2022), and it is unknown whether these ratios remain constant once sexual maturity is reached. Body mass and total length were not included in the analysis as both can be impacted by caudal autotomy, which is common in tokay geckos (Nurhidayat et al. 2020; pers. obs.). Morphological traits pertaining to the tail (e.g., scales between second and third tail tubercle rows; Rösler et al. 2011) were not utilised for the same reason. Pre-cloacal pores were not included in the analysis as their mean value differs by sex (Rösler 2005). In total, tokay geckos were analysed for 18 distinct morphological traits (Table 1; Appendices S1 and S3-S5). Some traits were bilateral, resulting in 28 morphological values for each specimen. Thirty-eight Florida tokay geckos were analysed, along with the four aforementioned native-range specimens.

**Table 1.** Morphological traits of *Gekko gecko* as analysed in this study.

| Morphological trait (abbreviation) | Description | Bilateral |
| --- | --- | --- |
| Snout-vent length (SVL) | The length in millimeters between the tip of the snout and the cloaca. | No |
| Nasals (N) | The number of scales in direct contact with the nostril, excluding supralabials. | Yes |
| Internasals (I) | The number of scales in direct contact with the rostral scale, excluding supralabials and nasals. | No |
| Prefrontals (PF) | The number of scales in direct contact with $\geq 1$ internasal, excluding the nasals and the rostral scale. | No |
| Supralabials (SPL) | The number of enlarged scales bordering the upper edge of the mouth, excluding the rostral scale. | Yes |
| Sublabials (SBL) | The number of enlarged scales bordering the lower edge of the mouth, excluding the mental scale. | Yes |
| Postmentals (PM) | The number of scales in direct contact with the mental scale, excluding the sublabials. | No |
| Parasublabials (PSL) | The number of scales in direct contact with the sublabials 1-6, excluding the mental scale, the postmentals, and sublabial seven, but including the gulars. | Yes |
| Gulars bordering the postmentals (GP) | The number of scales in direct contact with the postmentals, excluding the mental scale and the sublabials. | No |
| Interorbitals (IN) | The number of scales between the eyes, excluding the supraorbitals (aka supraoculars). This is measured as the lowest number of scales which form a continuous chain in a straight line between supraorbital eight (counted from the anterior towards the posterior) of each eye. The value is recorded as the mean of three valid counts. | No |
| Dorsal tubercle rows (DTR) | The number of tubercles located on the dorsum (back) counted between the lateral folds at the midline (defined as the line equidistant between the forelimb and hindlimb axilla) of the specimen's torso. The value is recorded as the mean of three valid counts. | No |
| Lateral tubercles (LT) | The number of tubercles in the ventral-most longitudinal row of tubercles on the flank. | Yes |
| Granules surrounding dorsal tubercles (GSDT) | The number of scales in direct contact with a randomly selected, large dorsal tubercle. The value is recorded as the mean of valid counts for three separate tubercles. | No |
| Thigh tubercles (TT) | The number of tubercles on the dorsal portion of the back leg, between the knee and hip. | Yes |
| Subdigital lamellae under the first finger (LF1) | The number of lamellae present on the 1<sup>st</sup> / inner-most digit of the forelimb. | Yes |
| Subdigital lamellae under the fourth finger (LF4) | The number of lamellae present on the 4<sup>th</sup> digit of the forelimb. | Yes |
| Subdigital lamellae under the first toe (LZ1) | The number of lamellae present on the 1<sup>st</sup> / inner-most digit of the hindlimb. | Yes |
| Subdigital lamellae under the fourth toe (LZ4) | The number of lamellae present on the 4<sup>th</sup> digit of the hindlimb. | Yes |

Discriminant Analysis of Principal Components (DAPC) (Jombart et al. 2010) was used to examine population-level morphological structure for the 38 Florida tokay geckos for which morphological data were available (“DAPC1”). DAPC was originally designed for genetic data, but is suitable for the analysis of morphological data (Jombart and Collins 2015), as evidenced in a recent study of the *G. smithii* species complex (Grismer et al. 2022). DAPC1 was performed on a morphological matrix using the *dapc* function in the ‘adegenet’ package (Jombart 2008) as executed in R version 4.0.5 (R Core Team 2022); the analysis was run using the clusters (*k*) returned by *find*.*clusters* (which performs a *k*-means clustering analysis in order to determine the *k* associated with the lowest Bayesian Information Criterion [BIC]; see Jombart and Collins 2015, Section 2.1), and retaining the number of Principal Components (PCs) associated with the lowest Root Mean Squared Error (RMSE) (as established using the *xvalDapc* function) (Jombart and Collins 2015). Additionally, the *predict*.*dapc* function was used to determine cluster membership probabilities for the four native-range tokay geckos. Thereafter, chi-squared tests were used to establish whether morphological cluster membership was non-randomly associated with mtDNA haplotypes (mtDNA Clade A or mtDNA Clade B), sex, or collection site (for sites with sample sizes ≥ 5). Student**’**s unpaired *t* tests (“*t* tests”) were used to check whether individual morphological traits differed between specimens possessing mtDNA Clade A haplotypes (“mtDNA Clade A specimens”) and those possessing mtDNA Clade B haplotypes (“mtDNA Clade B specimens”). α was set at 0.05 for all statistical tests.

A similar analysis (“DAPC2”) was then conducted using a second morphological matrix, comprising the four native-range specimens and 18 Florida tokay geckos for which both mito-nuclear (i.e., ND2 and RAG-1) and morphological data were available. *predict*.*dapc* was not used to determine probable cluster membership for the native-range tokay geckos due to their presence in the morphological matrix. The number of morphological clusters to be returned by *find*.*clusters* was also set to *k* = 2, to test whether a “*G. g. gecko*” cluster (including the Philippines specimens) and a “Form B” cluster (including the Thailand specimens) would be returned. Due to both the smaller sample size and the inclusion of native-range specimens in the morphological matrix, chi-squared tests were not performed to determine if genotype was non-randomly associated with Florida collection site.

## Results

Bayesian phylogenetic analysis of ND2 (1090 bp) (261/1090 variable sites; average overall pairwise sequence divergence [excluding *G. smithii*] 6.87 %) (Fig. 1; Appendix S6) resulted in three well-supported clades. The first clade corresponded to mtDNA Clade A *sensu* Saijuntha et al. (2019) (also see Wang et al. 2013, Fieldsend et al. 2021b) and contained 12 Florida tokay geckos, as well as specimens from Cambodia, Indonesia, peninsular Malaysia, and the two Philippines specimens (FMNH 236071 [TWF351] and FMNH 266245 [TWF361]). The remaining 33 Florida tokay geckos placed within the second clade, along with specimens from peninsular Thailand and Ayeyarwady, Myanmar; this clade was thus analogous to mtDNA Clade B *sensu* Saijuntha et al. (2019). The third clade comprised a single specimen (CAS 213628; ND2 GenBank accession number JN019053; Rösler et al. 2011) from Magway, Myanmar, and was thus identical to mtDNA Clade C from Saijuntha et al. (2019).

**Figure 1.**
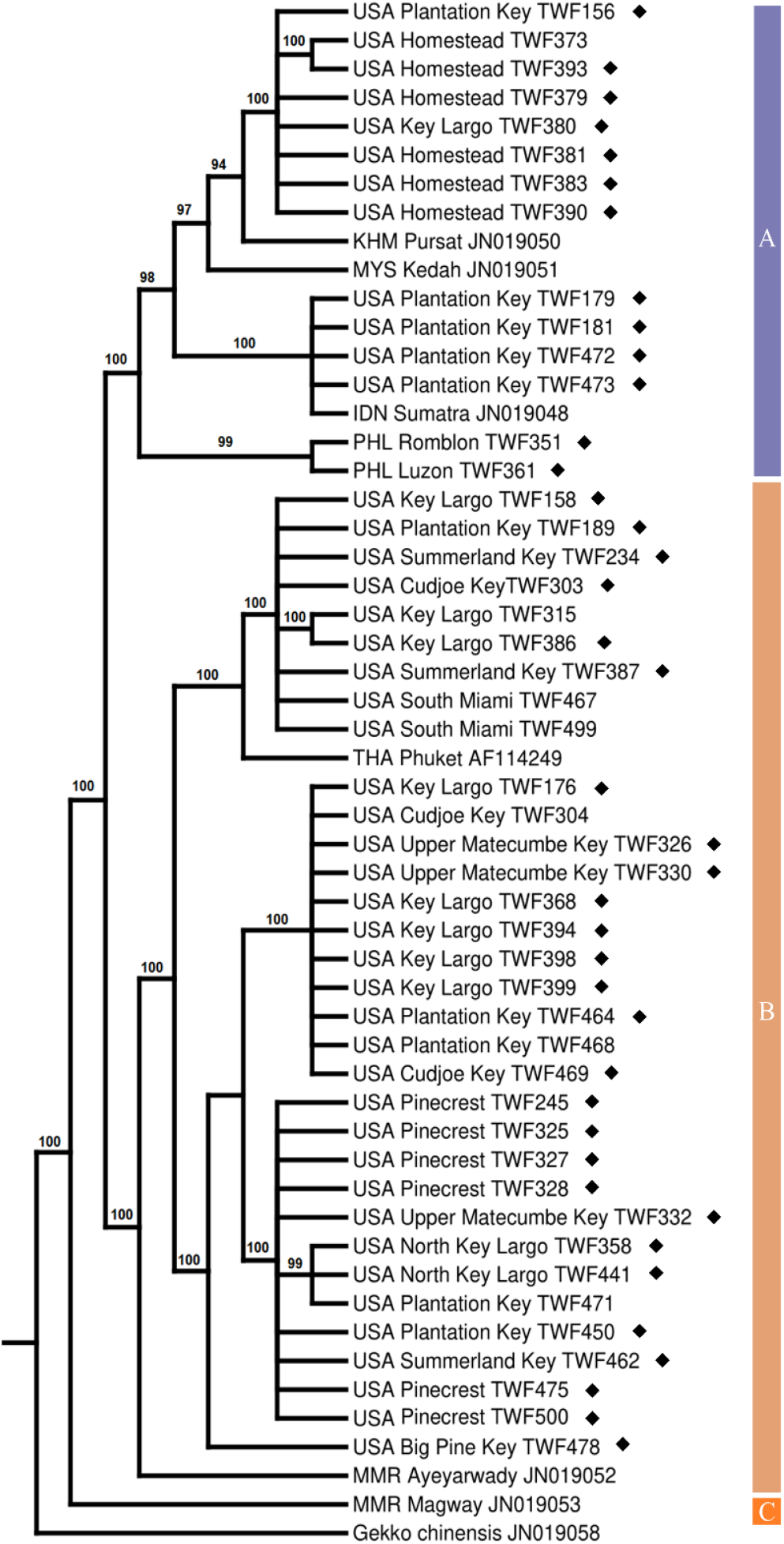
Bayesian 50 % majority-rule consensus tree (ND2, 1090 bp) for *Gekko gecko*. Numbers next to nodes indicate Bayesian posterior probabilities. Diamonds denote specimens that were analysed morphologically. Sequences generated as part of this study are identified by a unique ID number prefixed TWF (Appendix S1). GenBank accession numbers (two letters and six numerals) are given for sequences that were not generated as part of this study. Countries of origin are identified by their ISO 3166-1 alpha-3 codes: Cambodia – KHM; Indonesia – IDN; Malaysia – MYS; Myanmar – MMR; Thailand – THA; The Philippines – PHL; United States of America – USA. Clade names follow Saijuntha et al. (2019). Purple, light orange, and dark orange represent mtDNA Clades A, B, and C respectively.

Phylogenetic analysis of RAG-1 (553 bp) (10/553 variable sites; average over pairwise sequence divergence 0.41 %) (Fig. 2; Appendix S7) returned two major clades, with BPPs of 0.98 and 0.93. The first clade (nDNA Clade A) contained five Florida tokay geckos and the native-range specimens from Cambodia, Indonesia, peninsular Malaysia, and the Philippines. Twenty-one Florida tokay geckos clustered within the second clade (nDNA Clade B), alongside native-range specimens from peninsular Thailand and Ayeyarwady, Myanmar. The specimen from Magway, Myanmar, formed a sister clade to all other specimens in nDNA Clade A (BPP 0.99). While we refer to this sister clade as “nDNA Clade C” for consistency—i.e., to emphasise its relationship to mtDNA Clade C, and to distinguish it from the two nDNA lineages of primary interest in this study—it should be noted that in a more comprehensive RAG-1 gene tree (Appendix S8) this sequence formed part of nDNA Clade A proper, and did not form a sister clade (BPP 0.95).

**Figure 2.**
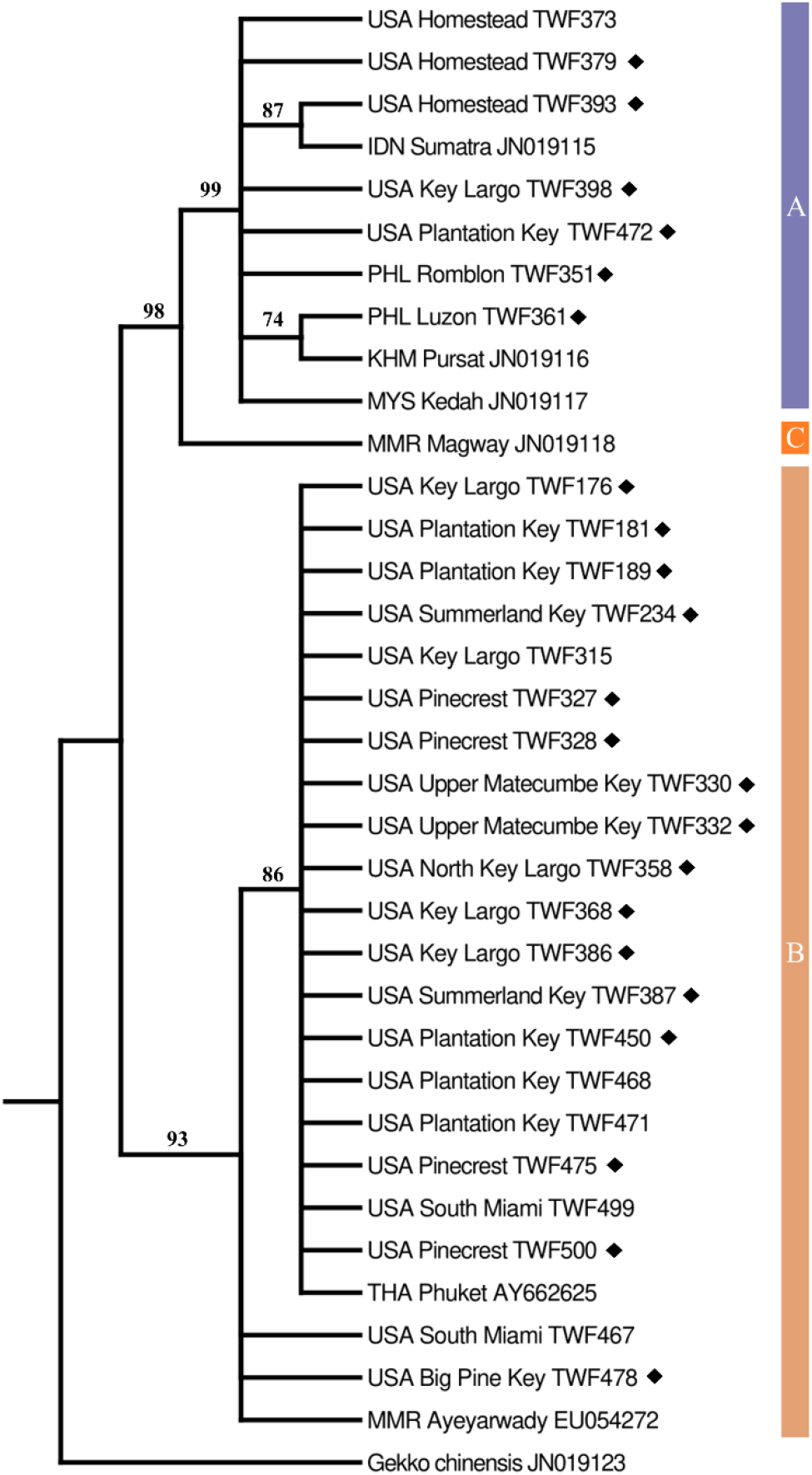
Bayesian 50 % majority-rule consensus tree (RAG-1, 553 bp) for *Gekko gecko*. Numbers next to nodes indicate Bayesian posterior probabilities. Diamonds denote specimens that were analysed morphologically. Sequences generated as part of this study are identified by a unique ID number prefixed TWF (Appendix S1). GenBank accession numbers (two letters and six numerals) are given for sequences that were not generated as part of this study. Countries of origin are identified by their ISO 3166-1 alpha-3 codes: Cambodia – KHM; Indonesia – IDN; Malaysia – MYS; Myanmar – MMR; Thailand – THA; The Philippines – PHL; United States of America – USA. Purple, light orange, and dark orange represent nDNA Clades A, B, and C respectively.

The concatenated ND2/RAG-1 dataset (1643 bp) (Fig. 3) produced a similar tree topology to the ND2 and RAG-1 phylogenies (Figs 1-2), with the Magway, Myanmar specimen forming its own distinct clade (“Lineage C”). The topology was also similar to the ND2 and RAG-1 phylogenies insofar as the Cambodia, Indonesia, peninsular Malaysia, and Philippines specimens from southeastern Asia clustered within the same clade (mtDNA Clade A or nDNA Clade A), and in a separate clade from the Ayeyarwady, Myanmar specimen (CAS 204952) (mtDNA Clade B or nDNA Clade B). The majority (21 of 26) of Florida tokay geckos clustered within the same clade as the Ayeyarwady specimen, while the remaining five clustered with the southeast Asian specimens. Of the 32 tokay geckos in this phylogeny not belonging to Lineage C, 10 (31.3 %) possessed mtDNA Clade A haplotypes and 10 (31.3 %) possessed nDNA Clade A alleles. Nine of the 10 specimens found to possess mtDNA Clade A haplotypes were also found to possess nDNA Clade A alleles (chi-squared test; *p* = < 0.001). Similarly, 21 of the 22 tokay geckos possessing mtDNA Clade B haplotypes were found to possess nDNA Clade B alleles (chi-squared test; *p* = 0.007). Thus, non-random association between mitochondrial and nuclear loci in this dataset was confirmed, and these two mito-nuclear lineages were named Lineage A (mtDNA Clade A co-occurring with nDNA Clade A) and Lineage B (mtDNA Clade B co-occurring with nDNA Clade B).

**Figure 3.**
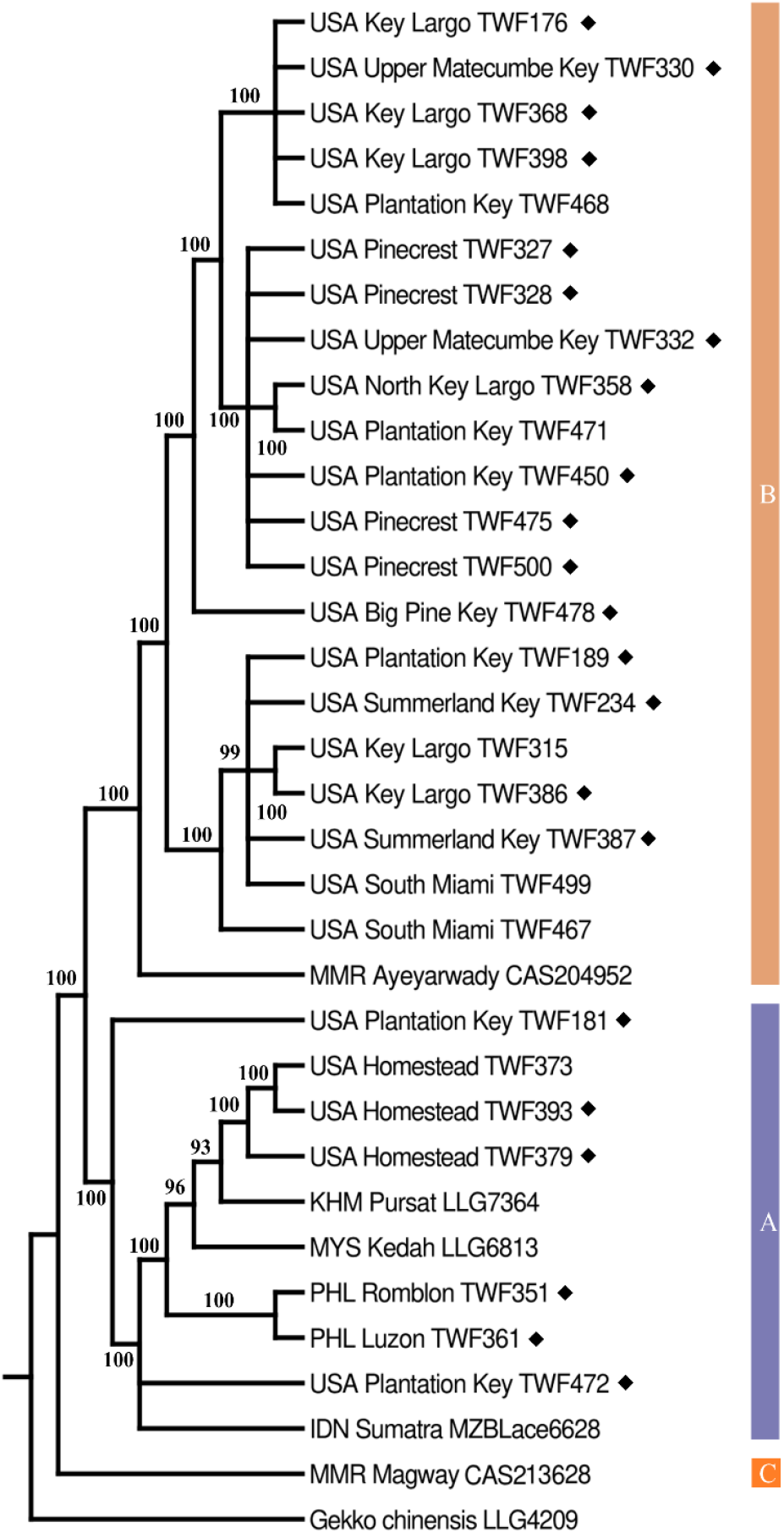
Bayesian 50 % majority-rule consensus tree (ND2, 1090 bp; RAG-1, 553 bp) for *Gekko gecko*. Numbers next to nodes indicate Bayesian posterior probabilities. Diamonds denote specimens that were analysed morphologically. Sequences generated as part of this study are identified by a unique ID number prefixed TWF (Appendix S1). Voucher numbers (as per Rösler et al. 2011) are given for sequences that were not generated as part of this study. Countries of origin are identified by their ISO 3166-1 alpha-3 codes: Cambodia – KHM; Indonesia – IDN; Malaysia – MYS; Myanmar – MMR; The Philippines – PHL; United States of America – USA. Purple, light orange, and dark orange represent Lineages A, B, and C respectively.

Collection site and mtDNA Clade affiliation for the 38 Florida tokay geckos analysed morphologically are shown in Fig. 4a. For DAPC1, *find*.*clusters* returned *k* = 2 as the number of morphological clusters with the lowest BIC. Six PCs—which together conserved 94.3% of the variance—were retained for DAPC1, along with one discriminant function. Twenty and 18 Florida tokay geckos were assigned to Clusters 1 and 2, respectively (Fig. 4b-d). Cluster 1 comprised two mtDNA Clade A specimens and 18 mtDNA Clade B specimens, while Cluster 2 contained nine mtDNA Clade A specimens and nine mtDNA Clade B specimens. Neither Cluster showed a biased sex ratio (Cluster 1, *p* = 0.494; Cluster 2, *p* = 0.471). Neither mtDNA Clade A nor mtDNA Clade B specimens were over-represented in Cluster 1 (*p* = 0.062), whereas mtDNA Clade A specimens were over-represented in Cluster 2 (*p* = 0.049). Specimens from Key Largo were over-represented in Cluster 1 (7 of 8, *p* = 0.049), while specimens from Plantation Key were over-represented in Cluster 2 (7 of 8, *p* = 0.023). No collection site was non-randomly associated with mtDNA Clade B specimens, but mtDNA Clade A specimens were over-represented at the Plantation Key and Homestead collection sites (Plantation Key 5 of 8, *p* = 0.036; Homestead 5 of 5; *p* = < 0.001). All four native-range tokay geckos were assigned to Cluster 2 by *predict*.*dapc* with high confidence (BPP ≥ 0.927).

**Figure 4.**
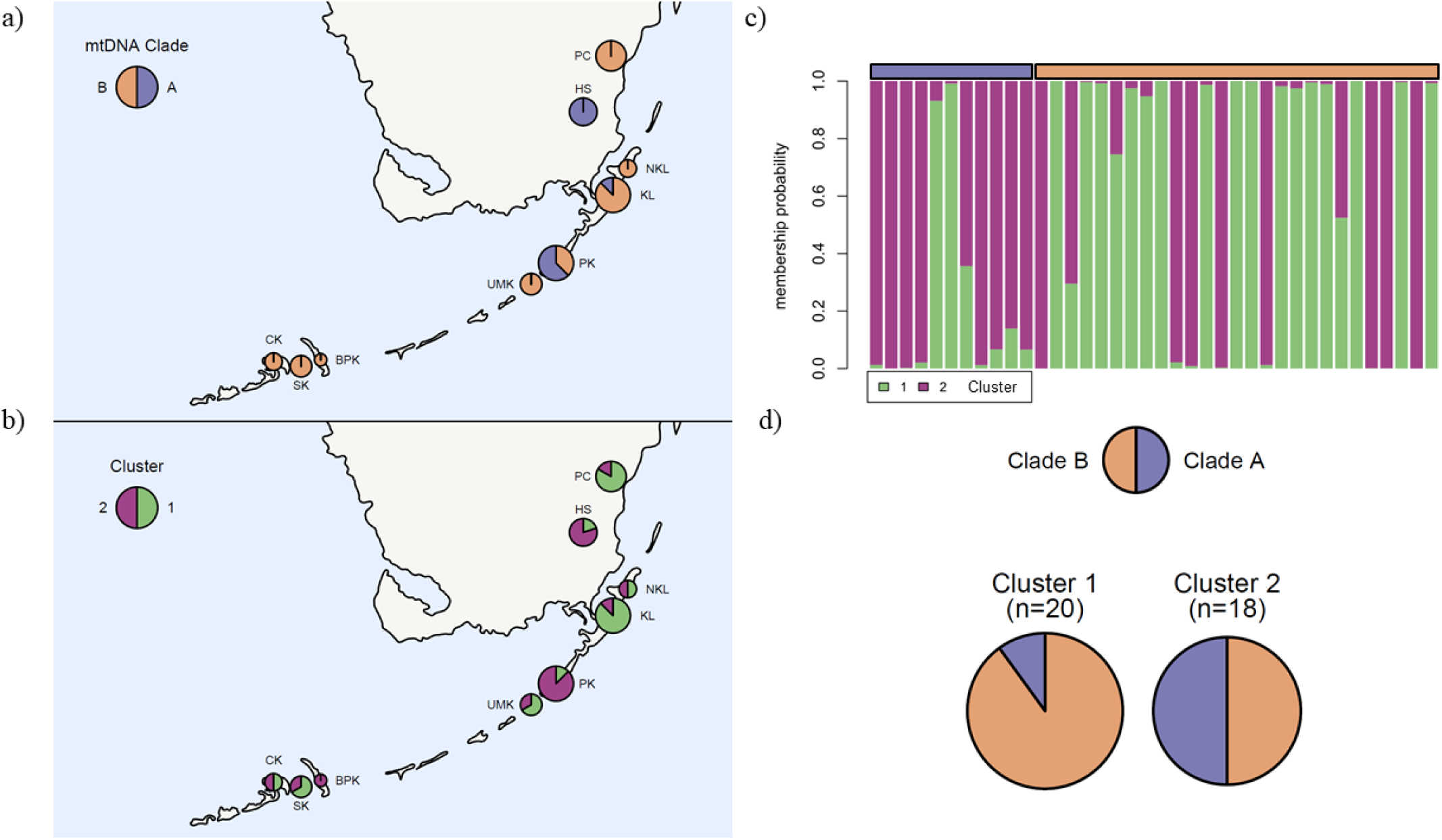
Locations and affiliations of Florida *Gekko gecko* (*n* = 38) included in this study by a) mtDNA clade and b) morphological cluster (*k* = 2) membership as determined by Discriminant Analysis of Principal Components (“DAPC1”). Map circle sizes correspond to population sample sizes (*n* = 1—8), with circle segment colouration representing the proportion of specimens from the population a) found to possess mtDNA Clade A vs mtDNA Clade B haplotypes or b) assigned to morphological Cluster 1 or 2. Site abbreviations are as follows: BPK – Big Pine Key; CK – Cudjoe Key; HS – Homestead; KL – Key Largo; NKL – North Key Largo; PC – Pinecrest; PK – Plantation Key; SK – Summerland Key; UMK – Upper Matecumbe Key. c) Morphological cluster membership probabilities; the purple and light orange horizontal bars denote specimens belonging to mtDNA Clades A and B, respectively. d) Pie charts showing the membership of the two morphological clusters by mtDNA clade.

The majority of morphological traits did not differ between mtDNA Clade A vs mtDNA Clade B specimens (*p* ≥ 0.05; Appendix S4), but mtDNA Clade B specimens were found to have longer SVLs, more lateral tubercles, and more thigh tubercles than mtDNA Clade A specimens (*p* ≤ 0.012; Appendix S4). Number of lateral tubercles or thigh tubercles did not strongly correlate with SVL (adjusted R^2^ values ≤ 0.13), confirming that the greater mean tubercle values observed in mtDNA Clade B specimens were not simply an artifact of their larger average body size.

As previously discussed, DAPC2 was performed on the four native-range tokay geckos and the 18 Florida tokay geckos for which mito-nuclear lineage and morphological data were available, and *k* was fixed at 2. DAPC2 was conducted using one discriminant function and three PCs, which together conserved 90.7% of the variance. Neither of the two clusters—named Cluster α and Cluster β to differentiate them from Clusters 1 and 2 (see above)—showed a biased sex ratio (Cluster α, *p* = 0.366; Cluster β, *p* = 0.257). Cluster α consisted of the two Thailand specimens that were not genotyped, and 11 Florida tokay geckos known to belong to Lineage B (“Lineage B specimens”). Cluster β contained four Lineage B specimens and all five tokay geckos belonging to Lineage A (“Lineage A specimens”), including the two Philippines specimens (Fig. 5). Cluster α was not associated with either Lineage (*p* = 0.055), but Lineage A was over-represented in Cluster β (*p* = 0.034). Most morphological traits were not significantly different between Lineage A vs Lineage B specimens (*p* ≥ 0.05; Appendix S5), but Lineage B specimens possessed fewer gulars bordering the postmentals, more interorbitals, more lateral tubercles on the left side of the body (viewed dorsally), and more thigh tubercles (both thighs) than Lineage A specimens *(p* ≤ 0.038; Appendix S5).

**Figure 5.**
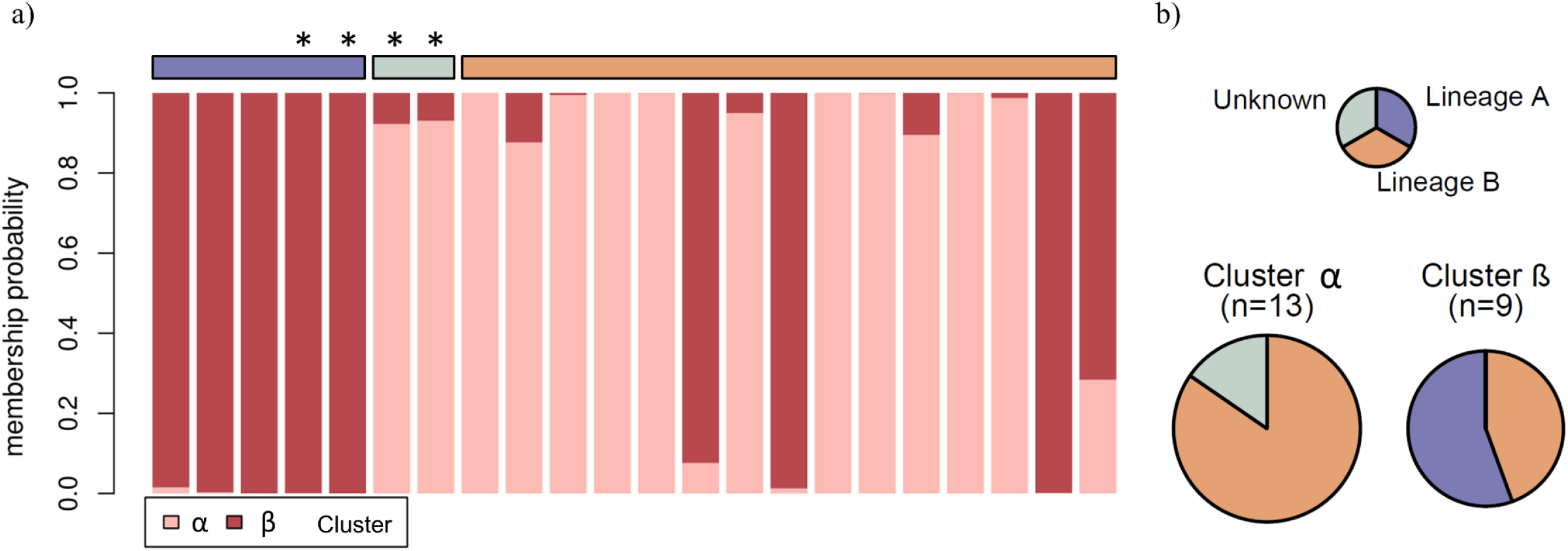
a) Morphological cluster (*k* = 2) membership probabilities for 18 Florida *Gekko gecko* and four native-range *G. gecko* as determined by Discriminant Analysis of Principal Components (“DAPC2”). The purple, light orange, and grey horizontal bars denote specimens belonging to Lineage A, Lineage B, and of unknown lineage respectively. Asterisks denote native-range specimens. b) Pie charts showing the membership of the two morphological clusters by Lineage.

## Discussion

This study used genotypic and phenotypic data from native-range and Florida tokay geckos to test the hypothesis of Fieldsend et al. (2021b) that multiple forms of the red-spotted tokay gecko—specifically, *G. g. gecko* and Form B—have been introduced to Florida, and to investigate the unresolved taxonomy of the red-spotted tokay gecko, *Gekko gecko*.

The ND2 mtDNA phylogeny accorded with the earlier findings of Fieldsend et al. (2021b) in identifying two mitochondrial DNA clades—mtDNA Clades A and B—in Florida tokay geckos (Fig. 1). Furthermore, phylogenetic analyses confirmed the existence of two distinct RAG-1 (nDNA) clades (Fig. 2; also see Appendix S8) in Florida. Importantly, mtDNA and nDNA clades were highly non-randomly associated (*p* ≤ 0.007) in this dataset, indicating the existence of two distinct mito-nuclear lineages in Florida, named Lineage A and Lineage B (see Results).

Reliable georeferenced native-range RAG-1 tokay gecko sequences are lacking (Fig. 2); however, analogous ND2 sequence data are far more abundant (Saijuntha et al. 2019, Fieldsend et al. 2021b). It is thus possible to state with confidence that the known distributions of mtDNA Clades A and B correspond closely with the putative distributions of *G. g. gecko* and Form B, respectively (Rösler 2005, Saijuntha et al. 2019, Fieldsend et al. 2021b). Given that mtDNA Clades A and B were found to be non-randomly associated with nDNA Clades A and B, respectively, it is highly probable that the native range distributions of mtDNA Clades A and B also reflect the native range distributions of the two distinct mito-nuclear lineages. i.e., Lineage A and Lineage B. This strongly suggests that Form B is not only differentiated from *G. g. gecko* phenotypically (Rösler 2005), but also genetically at the mito-nuclear level, and thus represents a distinct taxon. Form B may require elevation to species rank— perhaps under the available name *Gekko azhari*; however, a careful analysis of the literature and native-range specimens would first be required to ensure that such elevation was warranted.

Resolving the taxonomy of the tokay gecko *sensu lato* is of great importance, given the pressure placed on wild populations by collection for the Asian traditional medicine trade (Caillabet 2013, Nijman and Shepherd 2015, Weterings et al. 2019). For example, the native range of *G. g gecko* is currently characterised as spanning from Nepal to eastern Indonesia (Rösler et al. 2011), but the evidence presented here accords with the earlier assertion of Rösler (2005), i.e., that its range is actually restricted to the Malay Archipelago and relatively small areas of adjacent mainland southeastern Asia. Given that this region is at the epicentre of the global tokay gecko trade (Caillabet 2011, Caillabet 2013, Nijman and Shepherd 2015, Mallari 2019, Sy and Shepherd 2020), the threat posed to *G. g. gecko* by unsustainable harvesting may actually be much greater than is currently recognised. Furthermore, Fieldsend et al. (2021b) noted that undescribed forms of tokay gecko with highly restricted ranges may exist in Myanmar, a conjecture supported by the presence of Lineage C in Magway (Fig. 3); however, tokay geckos from Myanmar are probably illegally exported to China (Shepherd and Nijman 2007, Bauer 2009), thus illustrating how other unique forms of tokay gecko might also be threatened by the wildlife trade. A more complete picture of the phylogeography and taxonomy of the tokay gecko is thus crucial to ensuring that appropriate protections are put into place at the regional, national, and international levels.

Although DAPC1 (Fig. 4b-d) confirmed a link between genotype and phenotype in Florida tokay geckos—specifically, the non-random clustering of mtDNA Clade A specimens in morphological Cluster 2—two clear morphological forms representing *G. g. gecko* and Form B were not detected, an ambiguity most clearly illustrated by the assignation of all four native-range specimens to the same morphological cluster (Cluster 2). However, significant differences in the mean values of some traits— i.e., SVL and number of lateral and thigh tubercles—were still detected between Clade A and B specimens, providing further confirmation that genotype and phenotype are nevertheless linked to some degree in Florida tokay geckos, even when only a single mitochondrial gene is assayed.

DAPC2 (Fig. 5) was qualitatively similar to DAPC1, insofar as Lineage A specimens (all of which possessed mtDNA Clade A haplotypes) were non-randomly associated with a morphological cluster (in this case, Cluster β), whereas Lineage B specimens (all of which possessed mtDNA Clade B haplotypes) were not. However, unlike with DAPC1, DAPC2 assigned the native-range specimens to multiple clusters. Both Philippines specimens were assigned to Cluster β—which contained all five Lineage A specimens, and in which Lineage A predominated—while the Thailand specimens were assigned to Cluster α, which contained only Lineage B specimens. This result was consistent with what would be expected if Lineage A and Lineage B are indeed representative of *G. g. gecko* and Form B, respectively.

Snout-vent length data provide additional support for the key findings of this study. While Form B has a maximum SVL of 181 mm, the SVL of *G. g. gecko* does not exceed 162 mm (Rösler 2005). The five Florida tokay geckos with SVLs ≥ 162 mm genotyped in this study all possessed mtDNA Clade B ND2 haplotypes (Fig. 1; Appendix S1). RAG-1 was also sequenced for three of these specimens; all were found to possess nDNA Clade B alleles (Fig. 2), and to cluster within Lineage B (Fig. 3). Moreover, as Form B is the only form of tokay gecko whose maximum SVL exceeds 173 mm (Rösler 2005), the capture of a very large tokay gecko belonging to Lineage B on Key Largo (180 mm SVL; TWF386) essentially confirms the introduction of Form B to Florida, and further bolsters support for the link between morphological Form B and the Lineage B genotype.

One plausible explanation for the comparatively weak genotype-phenotype links observed in this study is the possible presence in Florida of hybrid or admixed tokay geckos with intermediate phenotypes. This phenomenon could also potentially explain the discrepancy in placement of native-range specimens by DAPC1 and DAPC2: as mtDNA is inherited matrilineally—and thus cannot be used as an indicator of hybridisation and admixture on its own (Avise et al. 1979)—the addition of a nDNA marker may have facilitated more-accurate genetic identification of specimens characteristic of one form of tokay gecko or the other. This would also explain why an equal number of morphological traits were found to differ significantly between Lineage A vs B specimens and mtDNA Clade A vs B specimens, despite the smaller sample size of the Lineage A vs B analysis (see Results). Different forms of tokay gecko probably hybridise in the wild (e.g, Wang et al. 2013), and Fieldsend et al. (2021b) speculated that they may do so in Florida. Hybridisation between *G. g. gecko* and Form B in Florida would itself be a concern, given that hybridisation can increase the invasive success of non-native organisms (e.g., Facon et al. 2005). Further study should thus be conducted to determine if hybridisation is occurring, and to establish whether it increases the risk posed to Florida’s native biodiversity by what may already be an invasive taxon (Fieldsend et al. 2021a, Fieldsend et al. 2021b).

In summary, our findings indicate both the existence of Form B as a distinct form of red-spotted tokay gecko, and the introduction of both *G. g. gecko* and Form B to Florida. Further study of populations in Florida and the native range is warranted in order to inform conservation efforts focussed on both the tokay gecko *sensu lato*, and Florida’s native biodiversity.

## Acknowledgements

We thank the Miami-Dade County Environmentally Endangered Lands Program and Miami-Dade County Parks, Recreation, and Open Spaces Department, Natural Areas Management Division; Castellow Hammock Preserve & Nature Center; Camp Owaissa Bauer; Florida Park Service; Florida Department of Environmental Protection Division of Recreation and Parks; United States Department of the Interior; United States Fish and Wildlife Service; FIU Tropics; the Susan S. Levine Trust; the FIU Institute of Environment; Coleman M. Sheehy III and Terry A. Lott of the Florida Museum of Natural History; Joshua Mata and Alan Resatar of the Field Museum of Natural History; Shahrzad Forouzanfar, Nathan Fox, Justin Bryan, Lilia Fernandez, Tyler Sanford, Paul Sharp, Alessandro Catenazzi, Joel Heinen, Sparkle Malone, Christina Romagosa, Rachunliu G. Kamei, and the public who generously assisted in various ways with the collection of specimens for this study. All work was performed under Florida International University IACUC protocol # IACUC-17-019.

## Funding

This work was supported by the FIU Institute of Environment; FIU Tropics; and the Susan S. Levine Trust.

## Notes

### Competing Interest Statement

The authors have declared no competing interest.

