## Appendix S3 for "Genotypic and phenotypic evidence indicates the introduction of two distinct forms of a non-native species (*Gekko gecko*) to Florida, USA"

### Appendix S3
### Morphological key for the tokay gecko

Thomas W. Fieldsend, Herbert Rösler, Kenneth L. Krysko, and Stephen Mahony

*This trait will not be directly comparable to specimens listed in Rösler et al. (2011). All other traits are included in the analyses of Rösler et al. (2011), although their exact definition may differ.

#### Glossary

*Lamella(e)* Subdigital scales with a width/height ratio exceeding 2:1 (sing. lamella; pl. lamellae).

*Parasublabial(s)* Synonym of Parainfralabial(s).

*Sublabial(s)* Synonym of Infralabial(s).

*Tubercle* a raised scale which is at least 1.5x the size of the adjacent scales. It is generally "wart-like" in appearance.

#### Annotation key

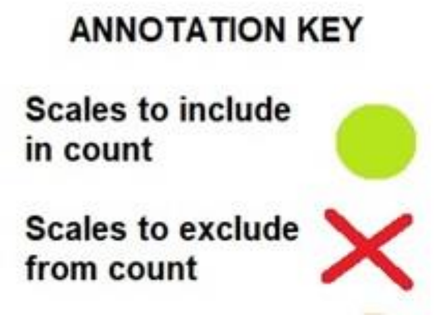

#### Sexing

Male (♂) tokay geckos tend to have pre-cloacal pores, post-cloacal sacs, and cloacal spurs that are larger/more pronounced that those of females (♀). Males are also larger than females on average, and generally have wider heads.

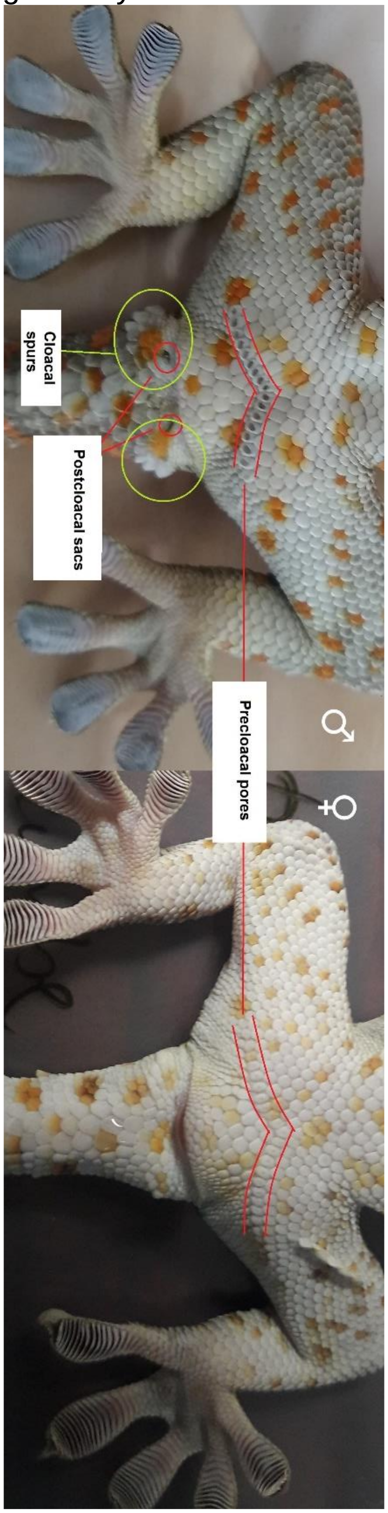

#### 1. Nasals (N)

The number of scales in direct contact with the nostril, excluding supralabials.

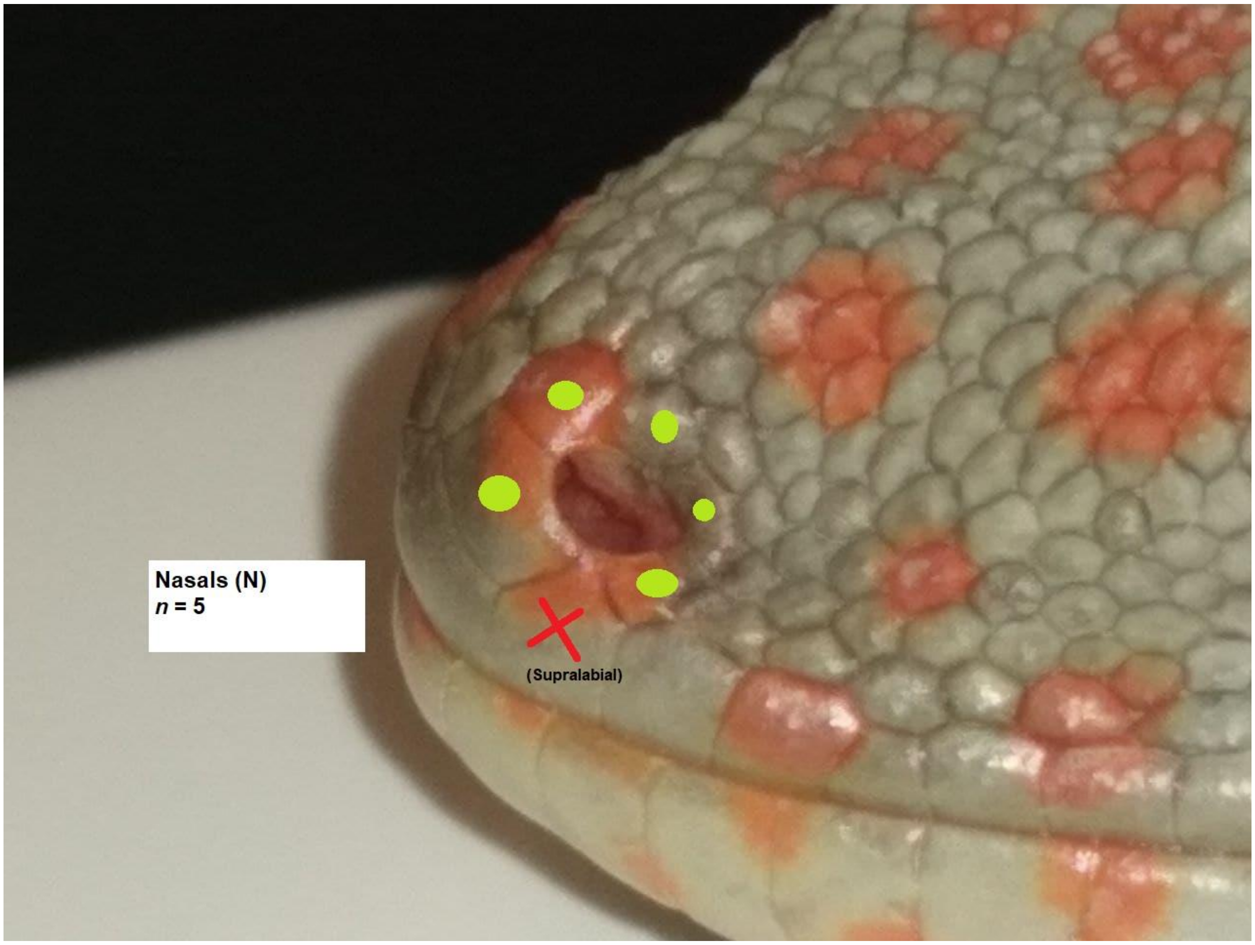

#### 2. Internasals (I)

The number of scales in direct contact with the rostral scale, excluding supralabials and nasals.

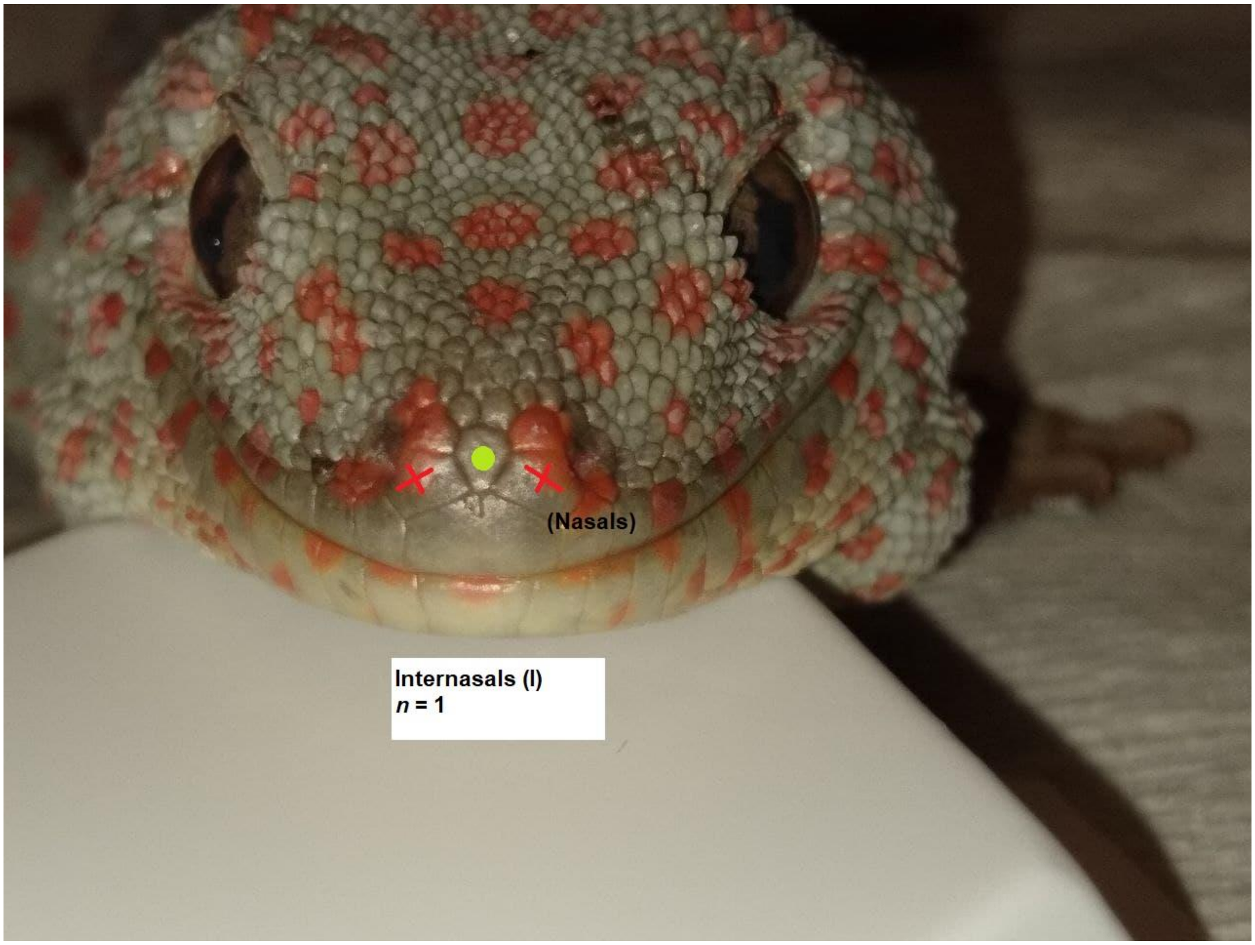

#### 3. Prefrontals (PF)*

The number of scales in direct contact with ≥ 1 internasal, excluding the nasals and the rostral scale.

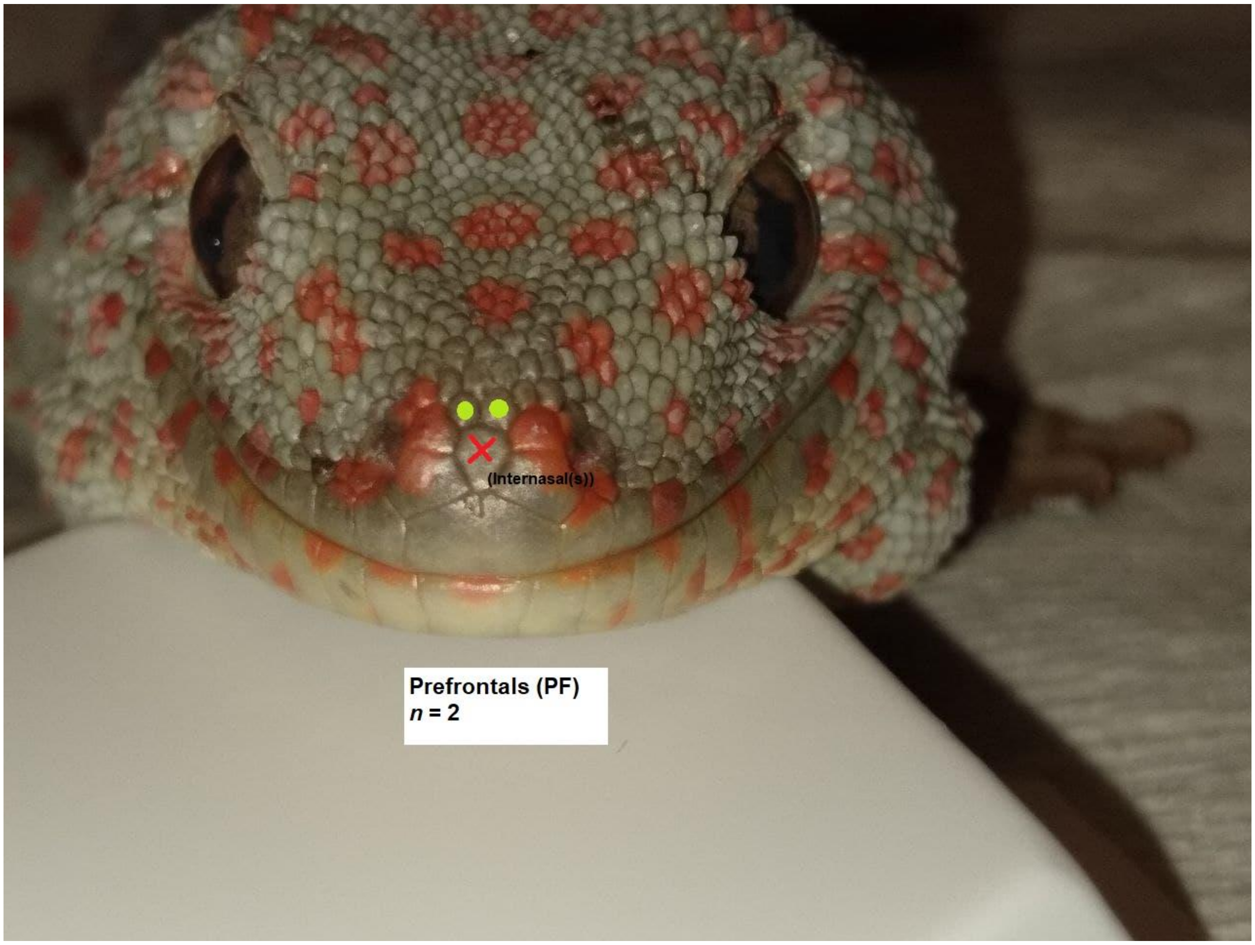

#### 4. Supralabials (SPL)

The number of enlarged scales bordering the upper edge of the mouth, excluding the rostral scale.

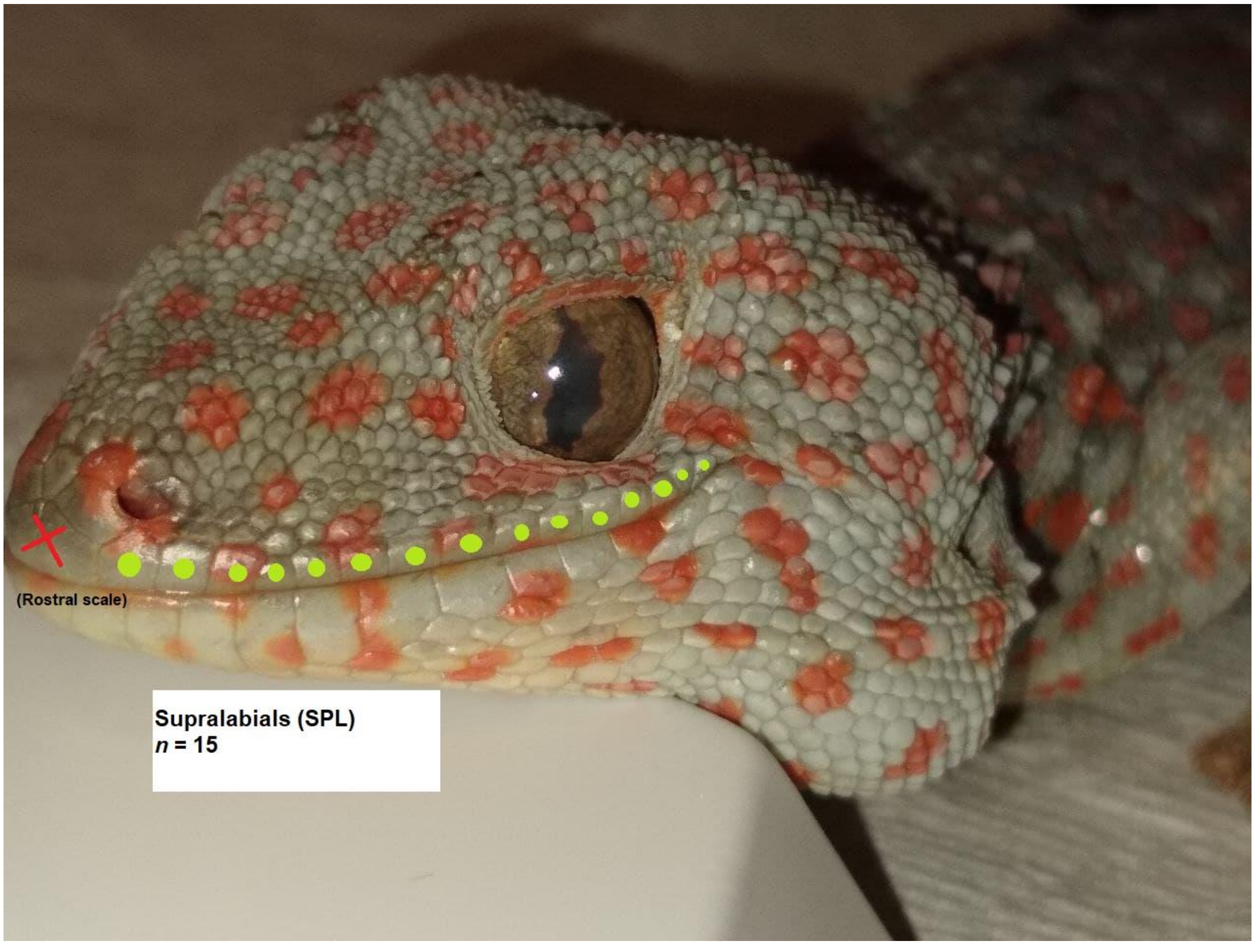

#### 5. Sublabials (SBL)

The number of enlarged scales bordering the lower edge of the mouth, excluding the mental scale.

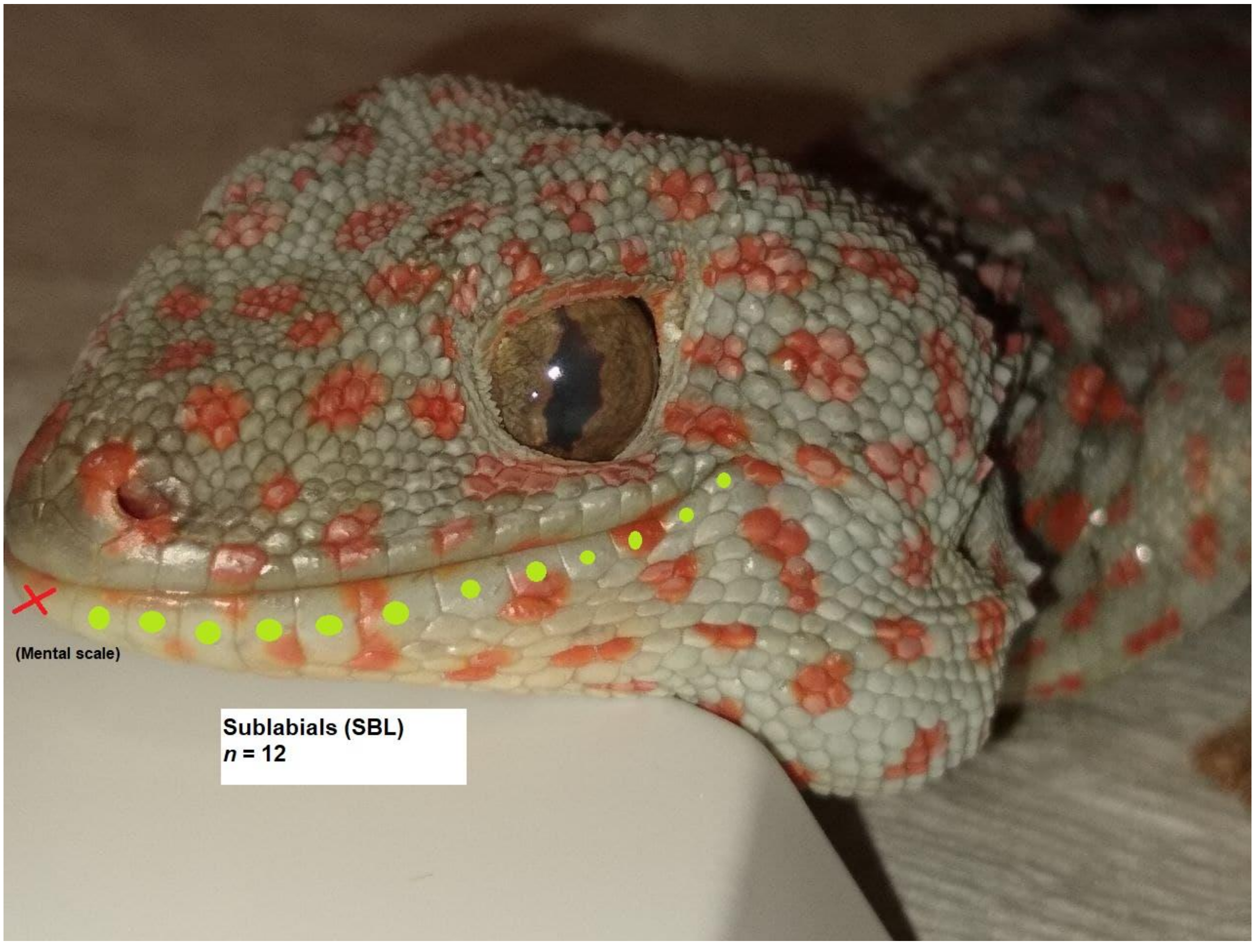

#### 6. Postmentals (PM)

The number of scales in direct contact with the mental scale, excluding the sublabials.

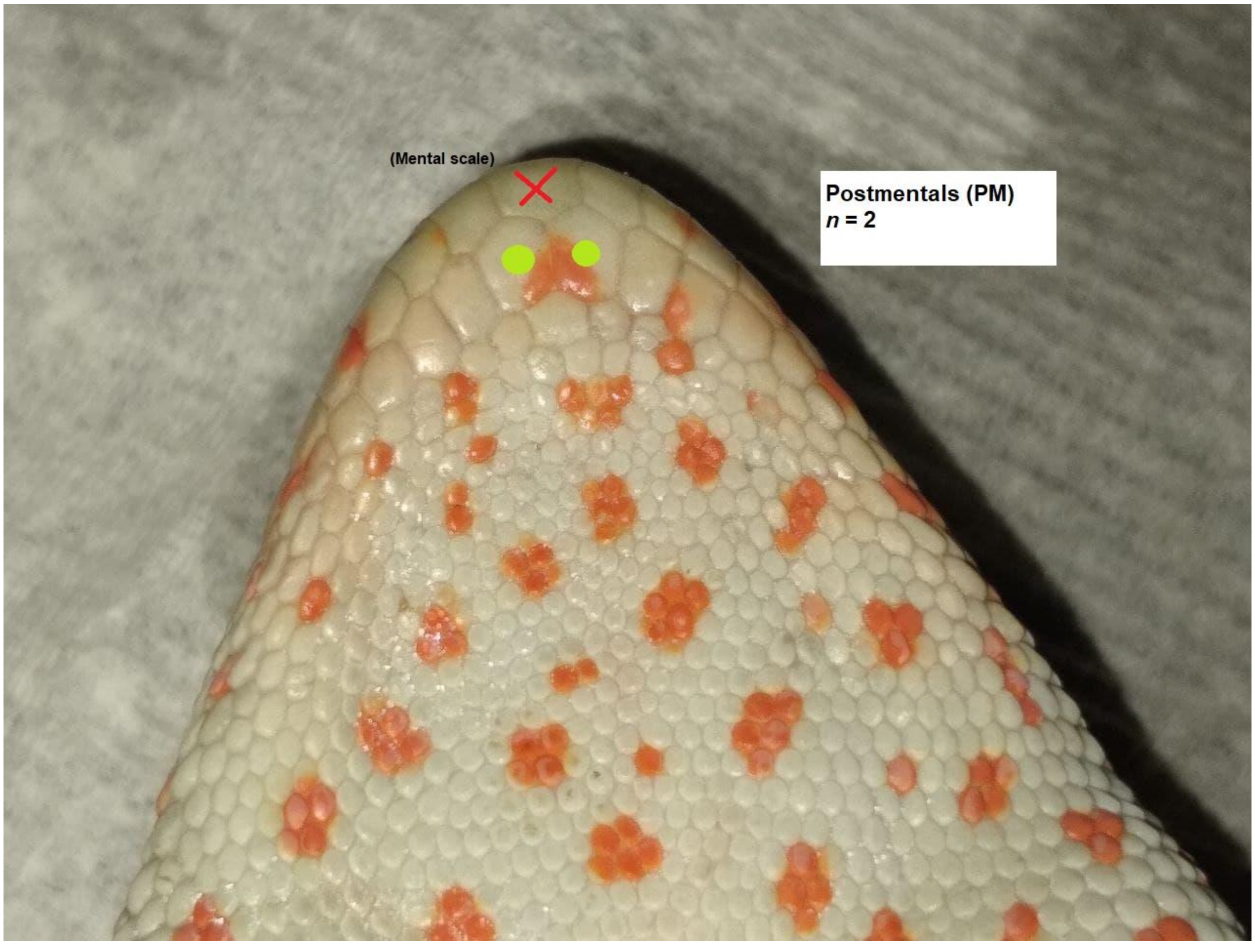

#### 7. Parasublabials (PSL)*

The number of scales in direct contact with the sublabials 1-6, excluding the mental scale, the postmentals, and sublabial seven, but including the gulars.

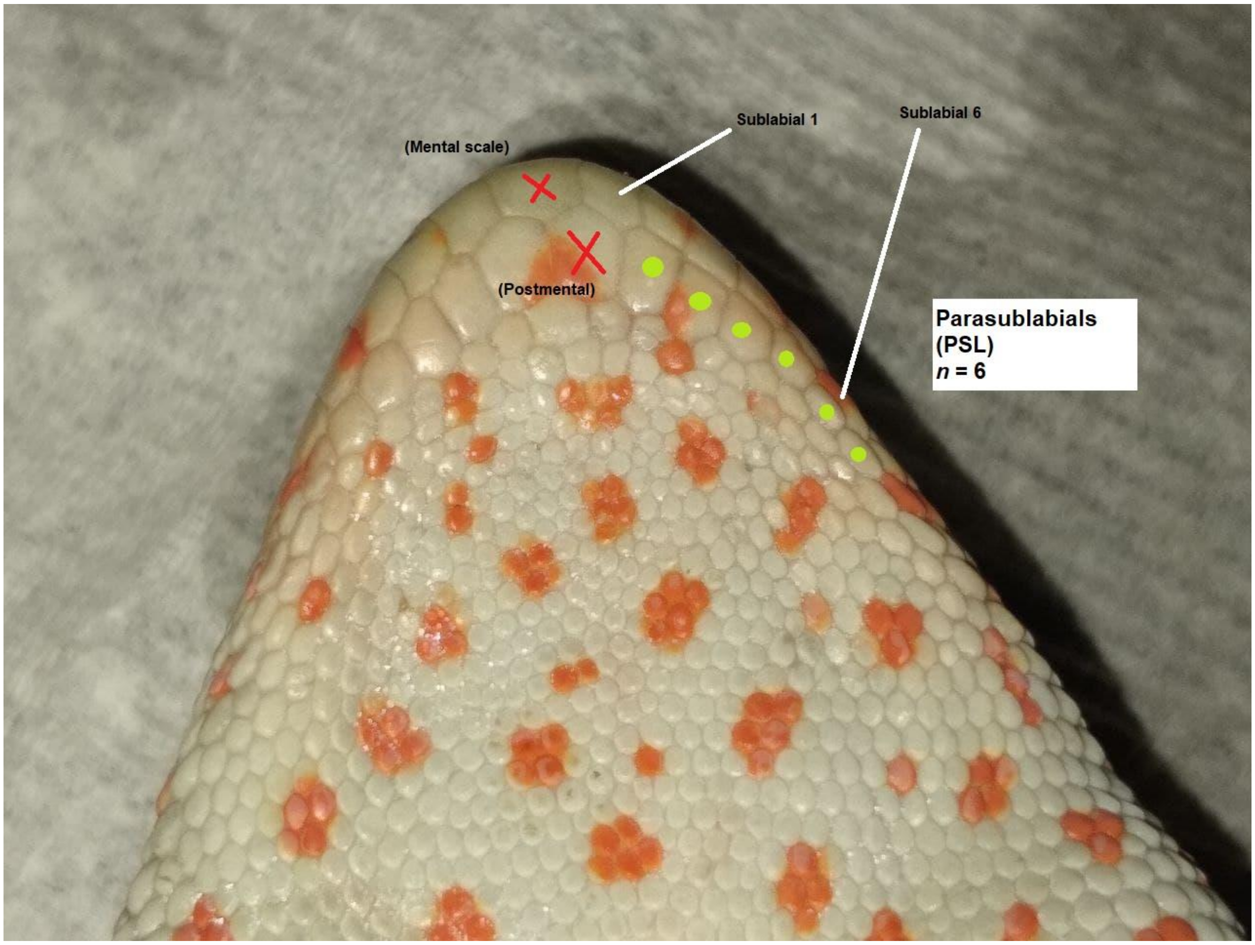

#### 8. Gulars bordering the postmentals (GP)

The number of scales in direct contact with the postmentals, excluding the mental scale and the sublabials.

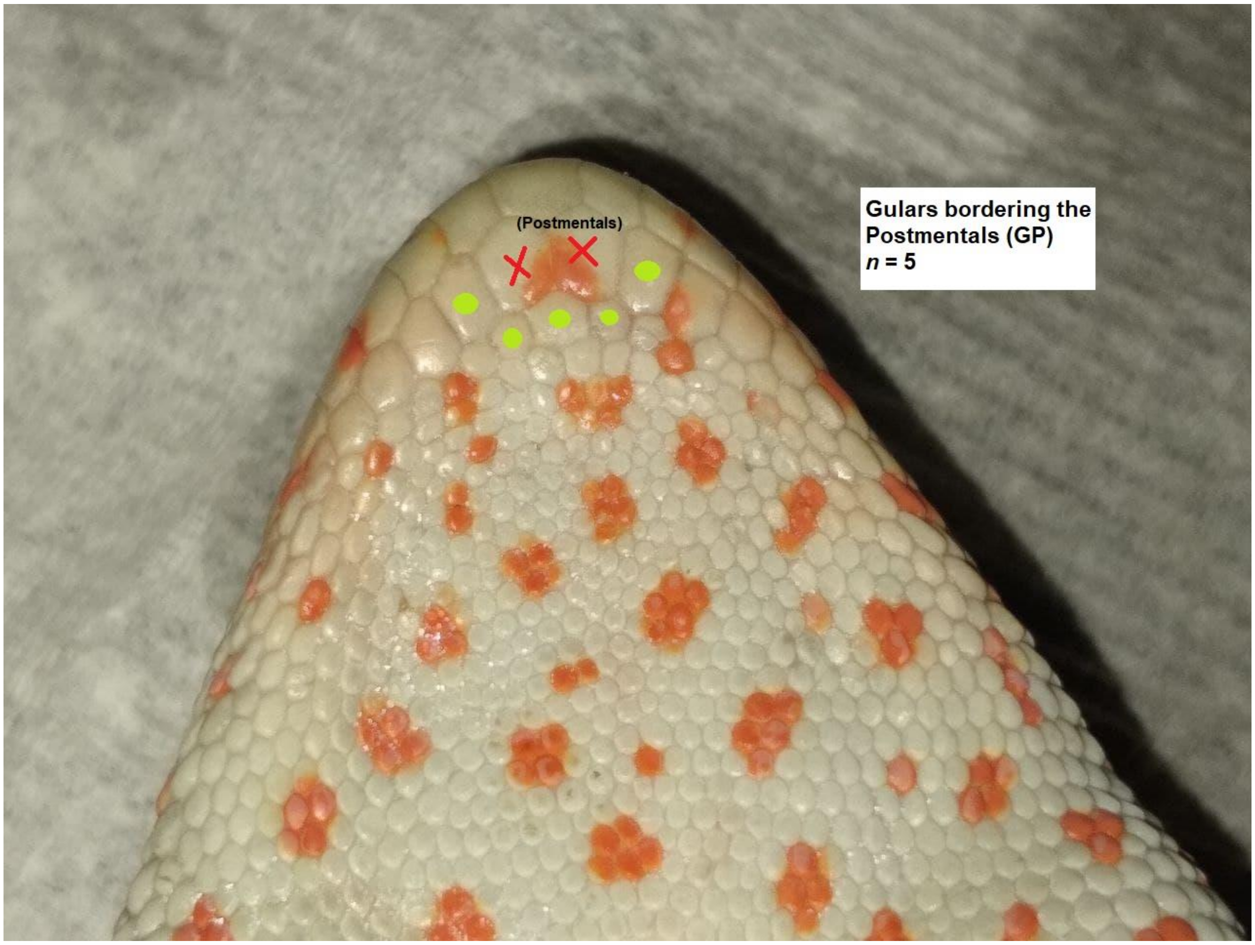

#### 9. Interorbitals (IN)

The number of scales between the eyes, excluding the supraorbitals (aka supraoculars). This is measured as the lowest number of scales which form a continuous chain in a straight line between supraorbital eight (counted from the anterior towards the posterior) of each eye. The value is recorded as the mean of three valid counts.

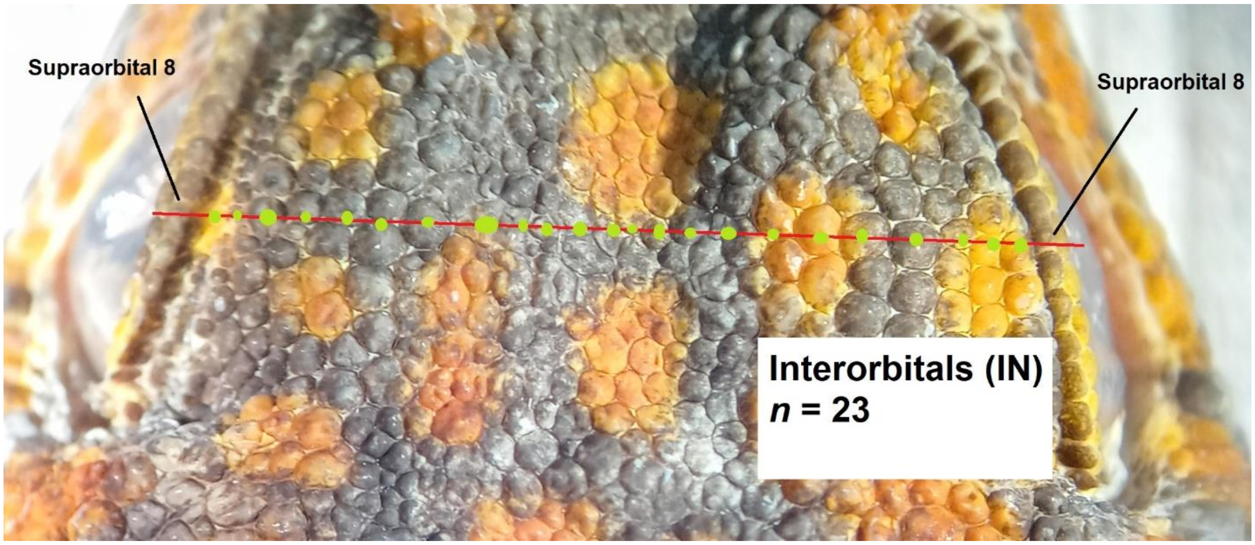

#### 10. Dorsal tubercle rows (DTR)

The number of tubercles located on the dorsum (back) counted between the lateral folds at the midline (defined as the line equidistant between the forelimb and hindlimb axilla) of the specimen's torso. The value is recorded as the mean of three valid counts.

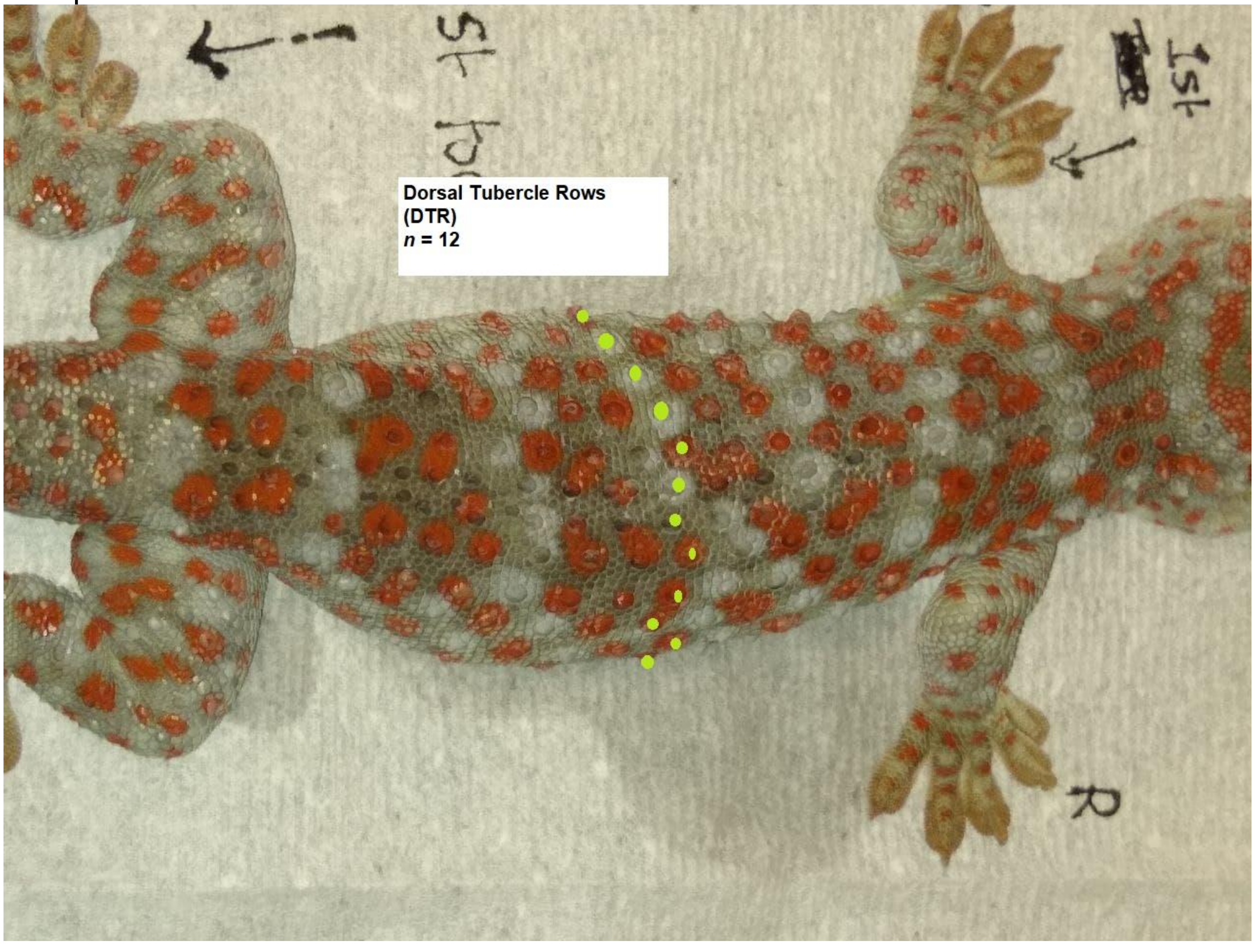

#### 11. Lateral tubercles (LT)*

The number of tubercles in the ventral-most longitudinal row of tubercles on the flank.

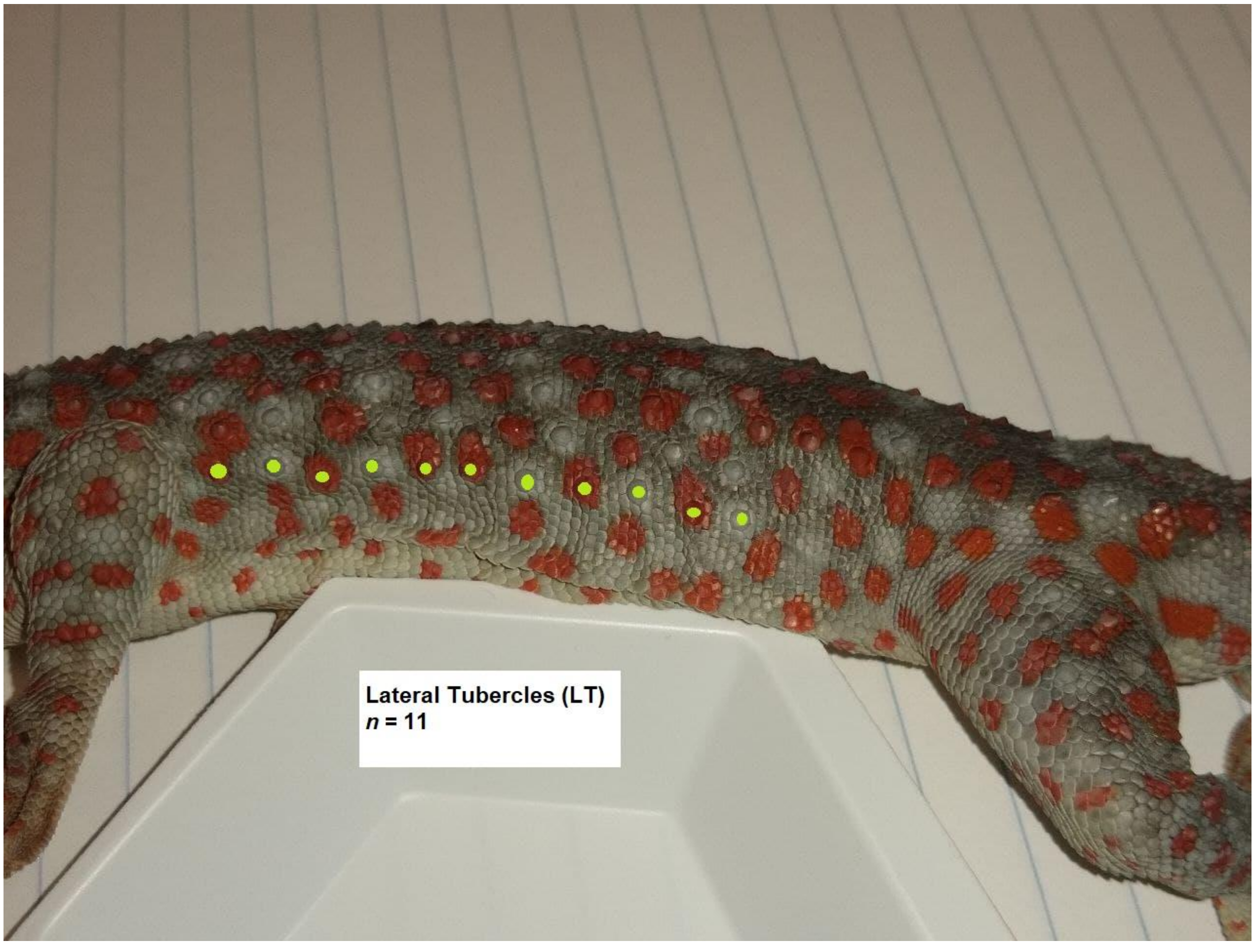

#### 12. Granules surrounding dorsal tubercles (GSDT)

The number of scales in direct contact with a randomly selected, large dorsal tubercle. The value is recorded as the mean of valid counts for three separate tubercles.

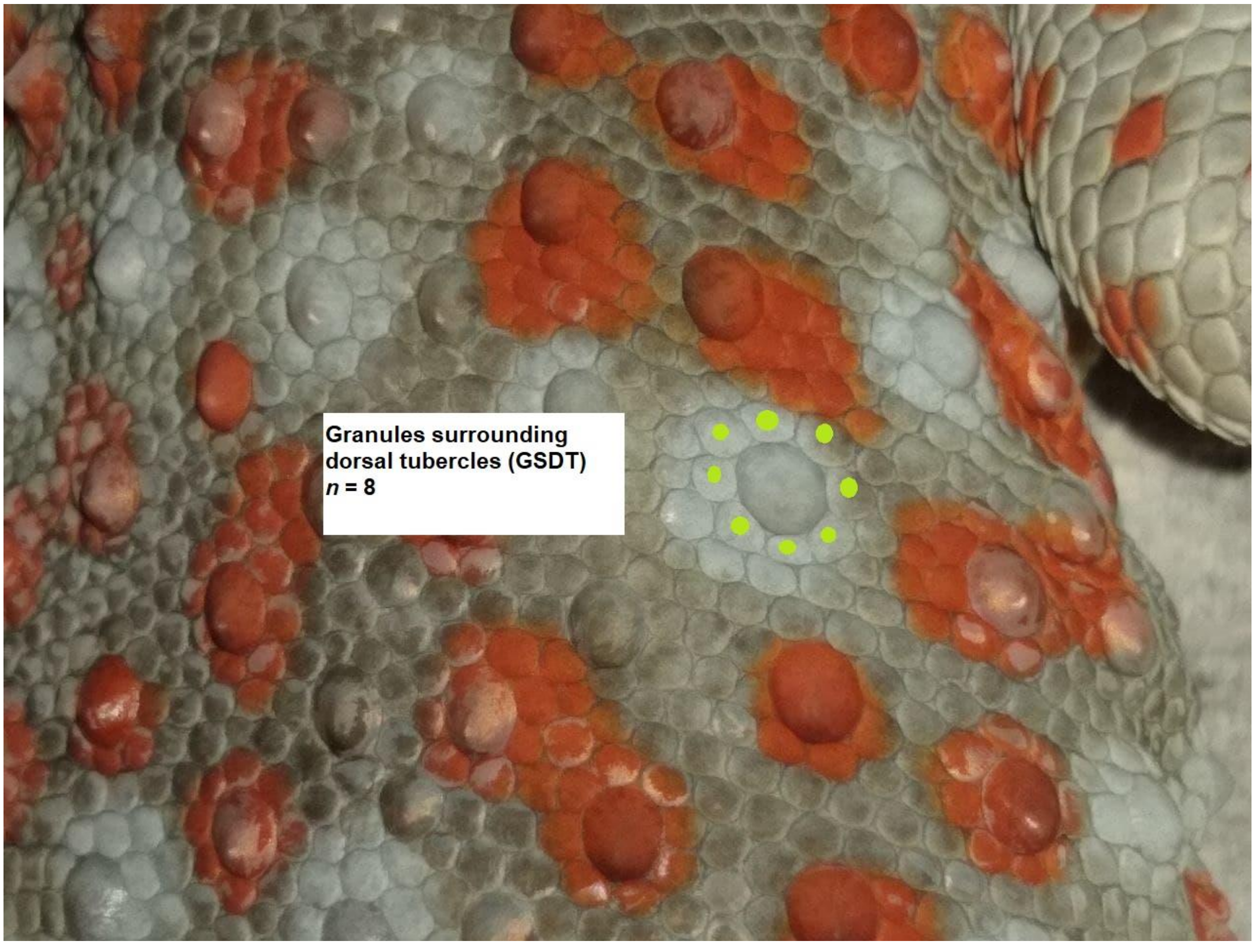

#### 13. Thigh tubercles (TT)

The number of tubercles on the dorsal portion of the back leg, between the knee and hip.

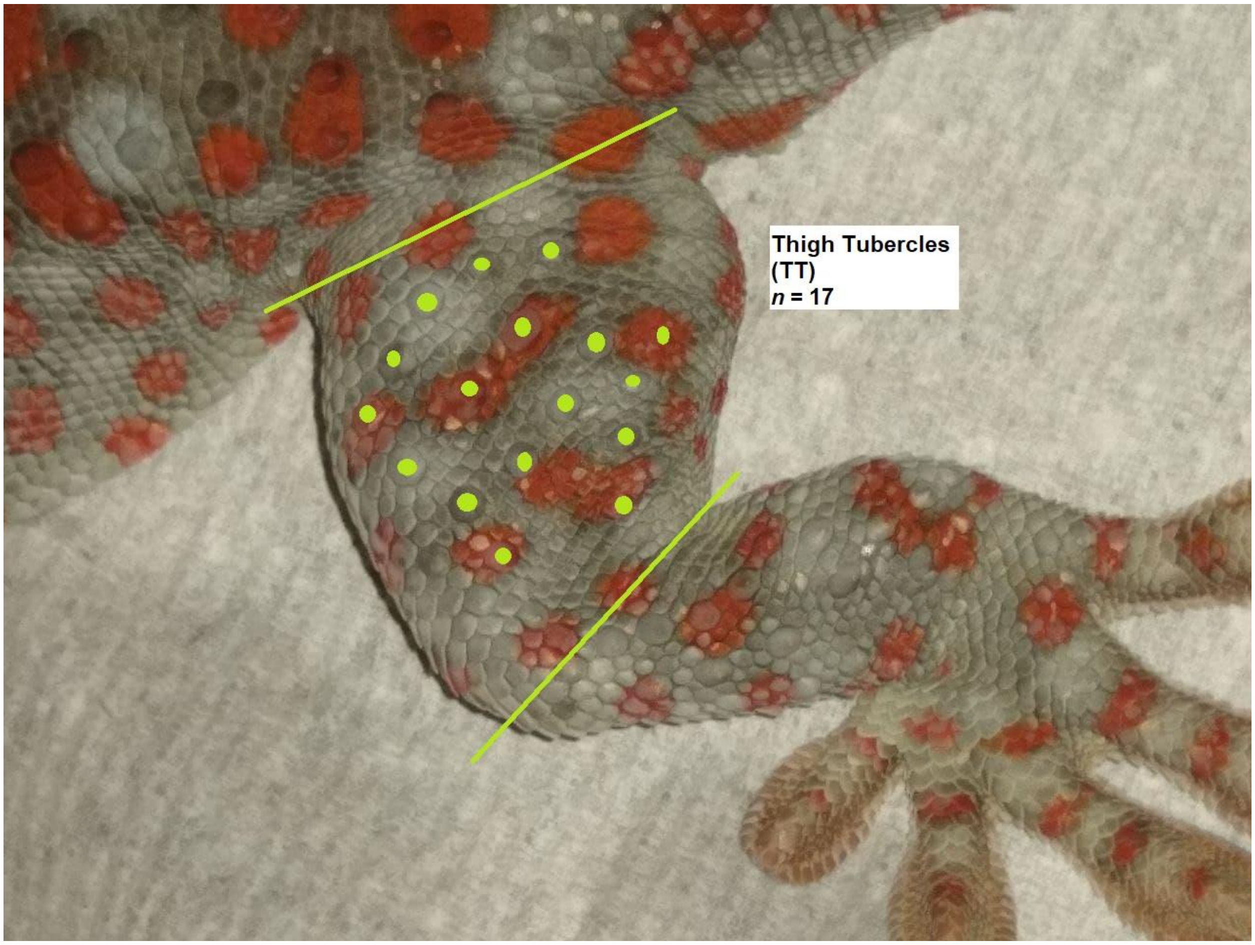

#### 14. Subdigital lamellae under the first finger (LF1)

The number of lamellae present on the 1st / inner-most digit of the forelimb.

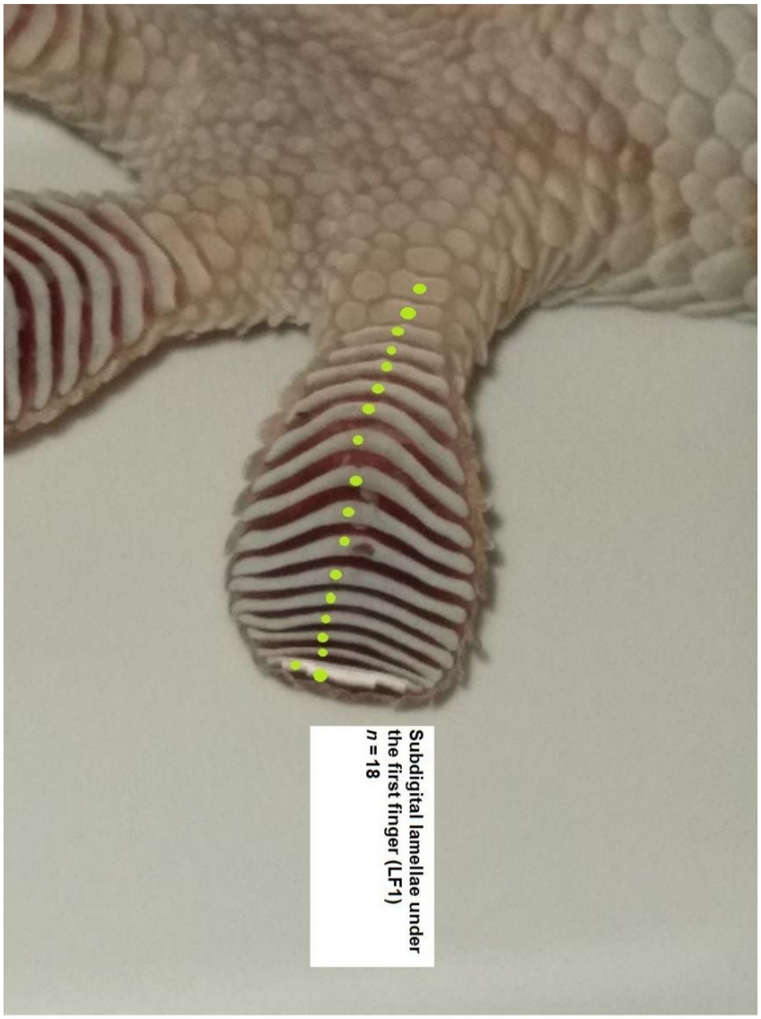

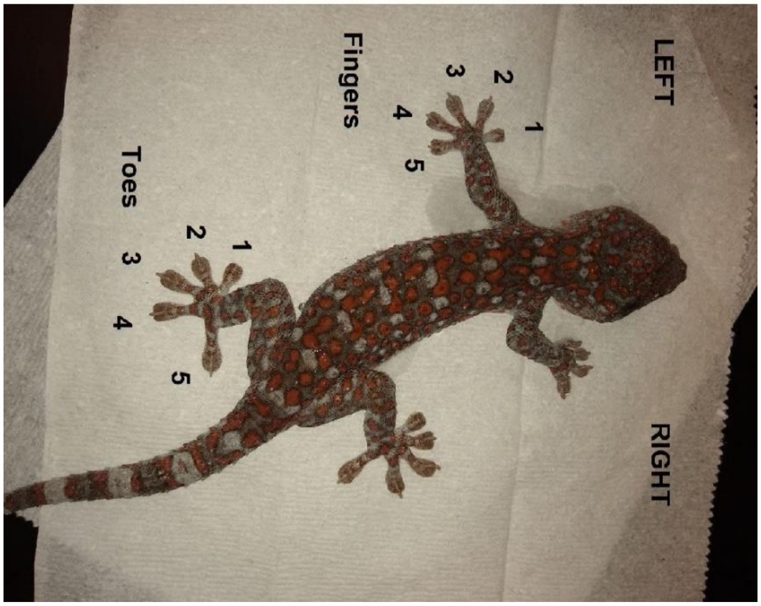

#### 15. Subdigital lamellae under the fourth finger (LF4)

The number of lamellae present on the 4th digit of the forelimb.

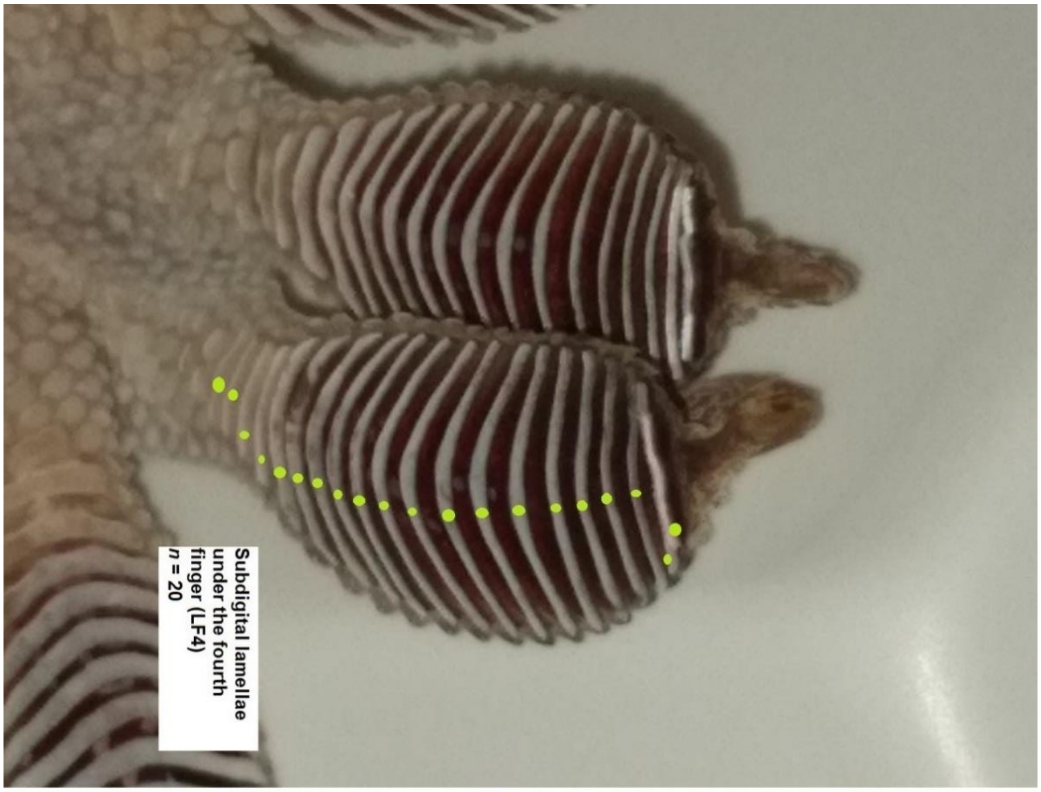

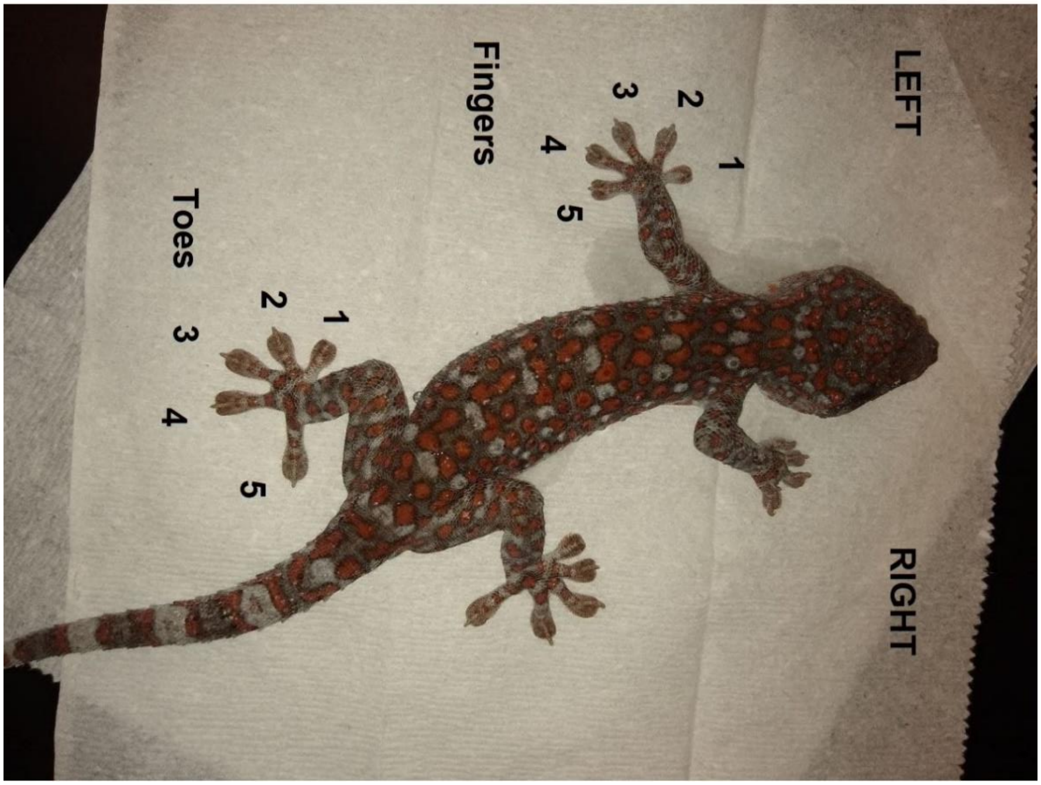

#### 16. Subdigital lamellae under the first toe (LZ1)

The number of lamellae present on the 1st / inner-most digit of the hindlimb.

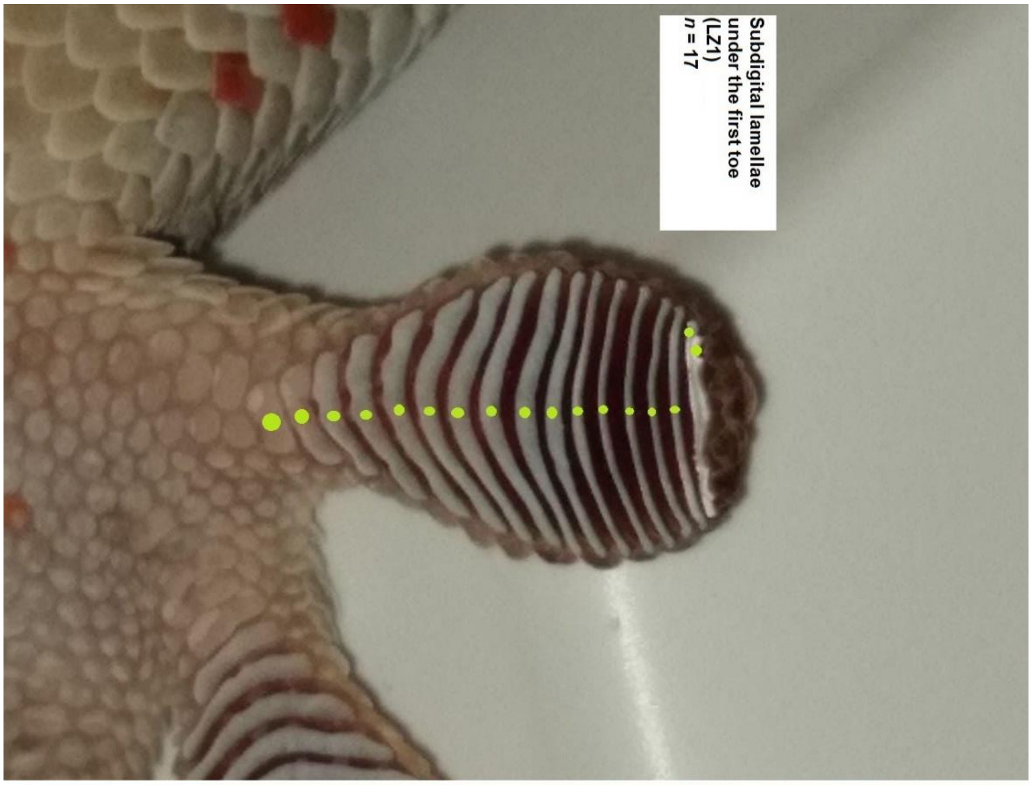

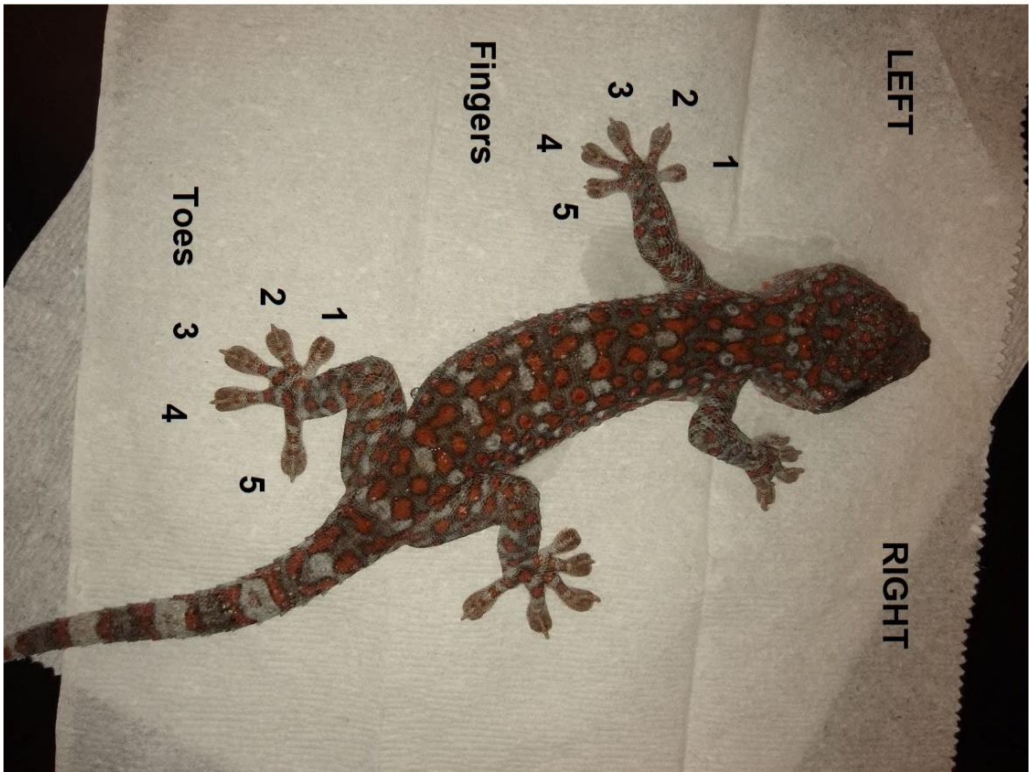

#### 17. Subdigital lamellae under the fourth toe (LZ4)

The number of lamellae present on the 4th digit of the hindlimb.

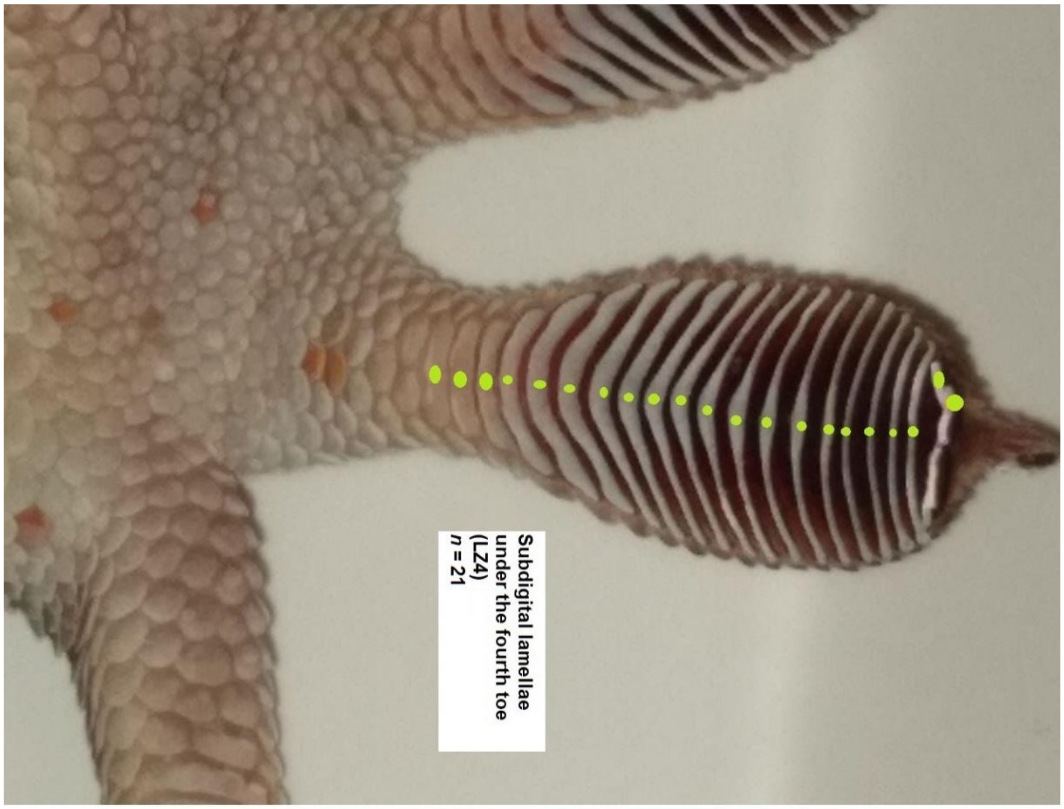

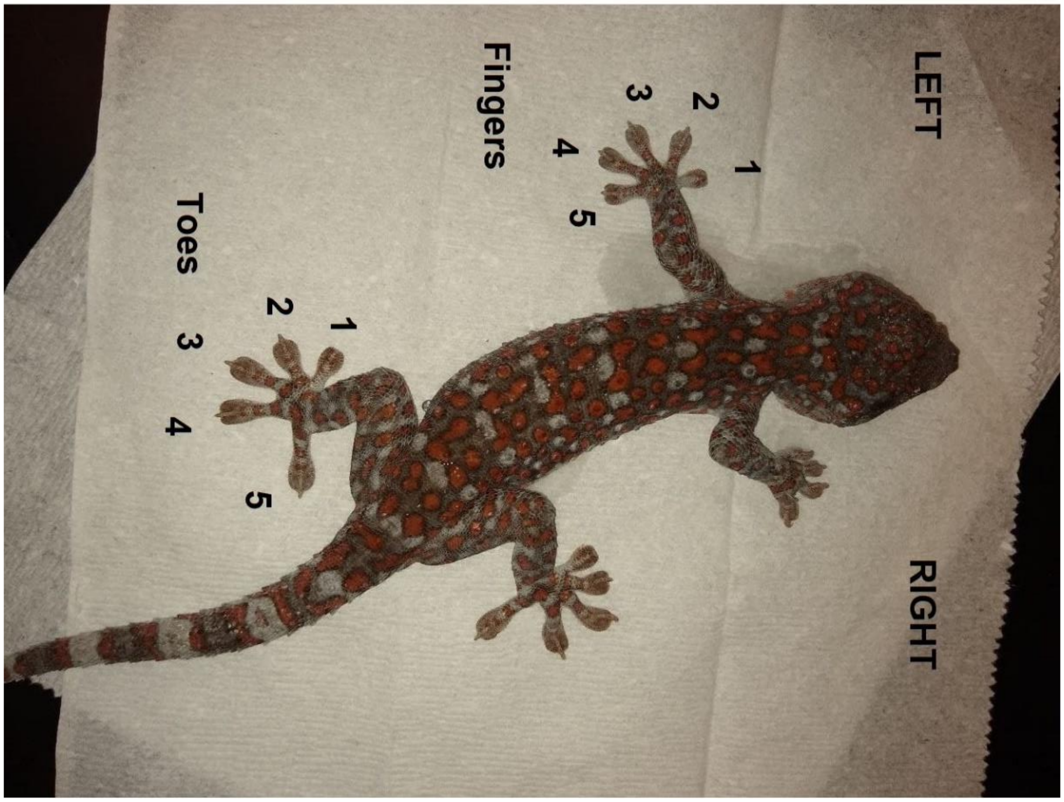
