## Appendix S8 for "Genotypic and phenotypic evidence indicates the introduction of two distinct forms of a non-native species (*Gekko gecko*) to Florida, USA"

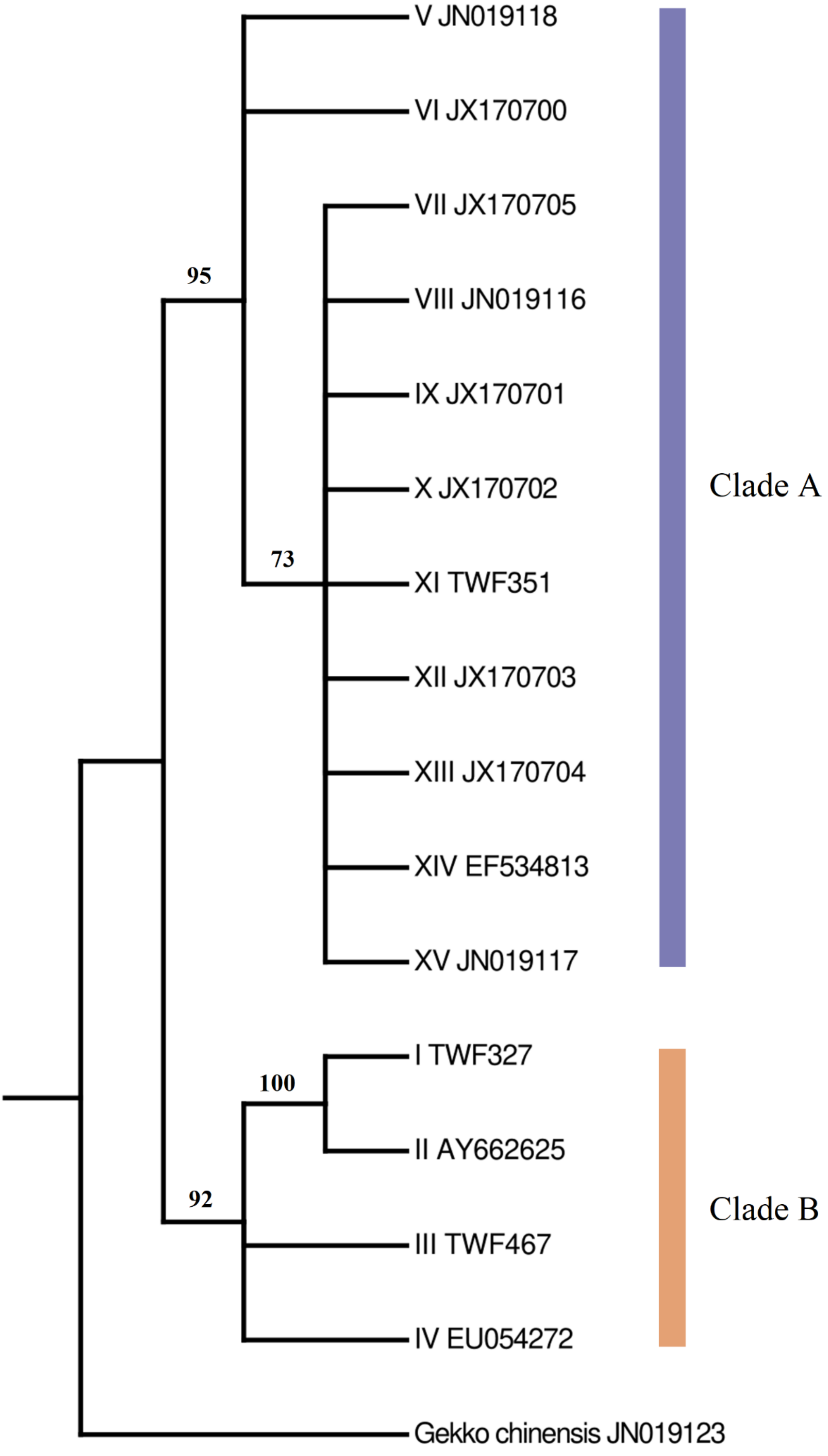
V JN019118
VI JX170700
VII JX170705
VIII JN019116
IX JX170701
X JX170702
XI TWF351
XII JX170703
XIII JX170704
XIV EF534813
XV JN019117
I TWF327
II AY662625
III TWF467
IV EU054272
Gekko chinensis JN019123
95
73
100
92
Clade A
Clade B

**Appendix S8** Bayesian 50 % majority-rule consensus tree (RAG-1, 546 bp) comprising all unique RAG-1 alleles currently recovered from the tokay gecko (*n* = 15). Refer to the manuscript proper for a description of the methods used to generate this gene tree. Numbers next to nodes indicate Bayesian posterior probabilities. Sequences generated as part of this study are identified by a unique ID number prefixed TWF; GenBank accession numbers (two letters and six numerals) are given for sequences that were not generated as part of this study. Sequences represent unique alleles rather than specimens and are thus named arbitrarily with Roman numerals. Purple and orange represent nDNA Clades A and B, respectively.
